# Ecological dynamics of the gut microbiome in response to dietary fiber

**DOI:** 10.1101/2021.08.20.457175

**Authors:** Hongbin Liu, Chen Liao, Jinhui Tang, Junyu Chen, Chaobi Lei, Linggang Zheng, Lu Wu, Chenhong Zhang, Yang-Yu Liu, Joao Xavier, Lei Dai

## Abstract

Dietary fibers are generally thought to benefit intestinal health. Their impacts on the composition and metabolic function of the gut microbiome, however, vary greatly across individuals. Previous research showed that each individual’s response to fibers depends on their baseline gut microbiome, but the ecology driving microbiota remodeling during fiber intake remained unclear. Here, we studied the long-term dynamics of gut microbiome and short-chain fatty acids (SCFAs) in isogenic mice with distinct microbiota baselines fed with the fermentable fiber inulin compared to the non-fermentable fiber cellulose. We found that inulin produced generally rapid response followed by gradual stabilization to new equilibria, and those dynamics were baseline-dependent. We parameterized an ecology model from the timeseries data, which revealed a group of bacteria whose growth significantly increases in response to inulin. and whose baseline abundance and interspecies competition explains the baseline-dependence of microbiome density and community composition dynamics. Fecal levels of of SCFAs, such as propionate, is associated with the abundance of inulin responders, yet inter-individual variation of gut microbiome impedes the prediction of SCFAs by machine learning models. Finally, we showed that our methods and major findings are generalizable to dietary resistant starch. This study emphasizes the importance of ecological modeling to understand microbiome responses to dietary changes and the need for personalized interventions.

## Introduction

Fermentable dietary fibers, such as inulin and resistant starch, are edible carbohydrate polymers that escape digestion by host enzymes in the upper gut and are fermented by gut microbiota in the cecum and colon. The major products from the microbial fermentative activity in the large intestine are short-chain fatty acids (SCFAs)—mainly acetate, propionate and butyrate—which have broad impacts on intestinal health and immunity [1-3]. Impaired SCFAs production has been associated with multiple dieseases [4, 5]. Dietary fibers can selectively enrich beneficial gut bacteria [6] and could be administered as “pre-biotic” therapies to restore intestinal gut microbiota and elevate SCFA levels [7, 8].

Previous work showed that dietary fibers can cause rapid changes in microbiota composition and biomass [9, 10]. But the ability of fibers to increase SCFA production varies across individuals [11-14]. For example, Baxter *et al*. showed that resistant starch was able to promote butyrate production in only 63% participants [12]. The individualized response can arise from a combination of factors such as host genetics and diet history. But the baseline gut microbiota is also a critical factor [14, 15]. The feces of some healthy human donors fail to ferment resistant starch, which can be restored by co-incubation with *R. brommi*, a well-known degrader of resistant starch [16]. Person-to-person variation in the bacterial and metabolomic composition of gut microbiome [17] can further impact biological variables such as body mass index [18] and glucose tolerance [19] of the human host.

Dietary fibers select from the pool of baseline community for microbial taxa that can use fibers as substrates for growth, and these responders could further impact the entire gut microbial community through a complex ecological network [20] (**Fig. 1A**). The primary users of dietary fibers are relatively few in low-fiber diet, but they can rapidly expand and dominate the gut microbiota after substantial induction [21]. Production of some SCFAs, especially butyrate, involves cross-feeding cooperations among specialized gut bacteria. By hydrolyzing complex polysaccharide fibers, primary degraders release into the gut partially breakdown products (e.g., mono- and oligo-saccharides) and fermentation metabolites (e.g., pyruvate), which can respectively benefit the secondary fiber degraders and SCFAs producers [22, 23]. Despite increasing interests in microbiota ecology and targeted modulation [24-27], a system-level, quantitative understanding of the ecological dyanmics of gut microbiome under dietary interventions and the dependence on the baseline community composition is still lacking.

**Figure 1.**
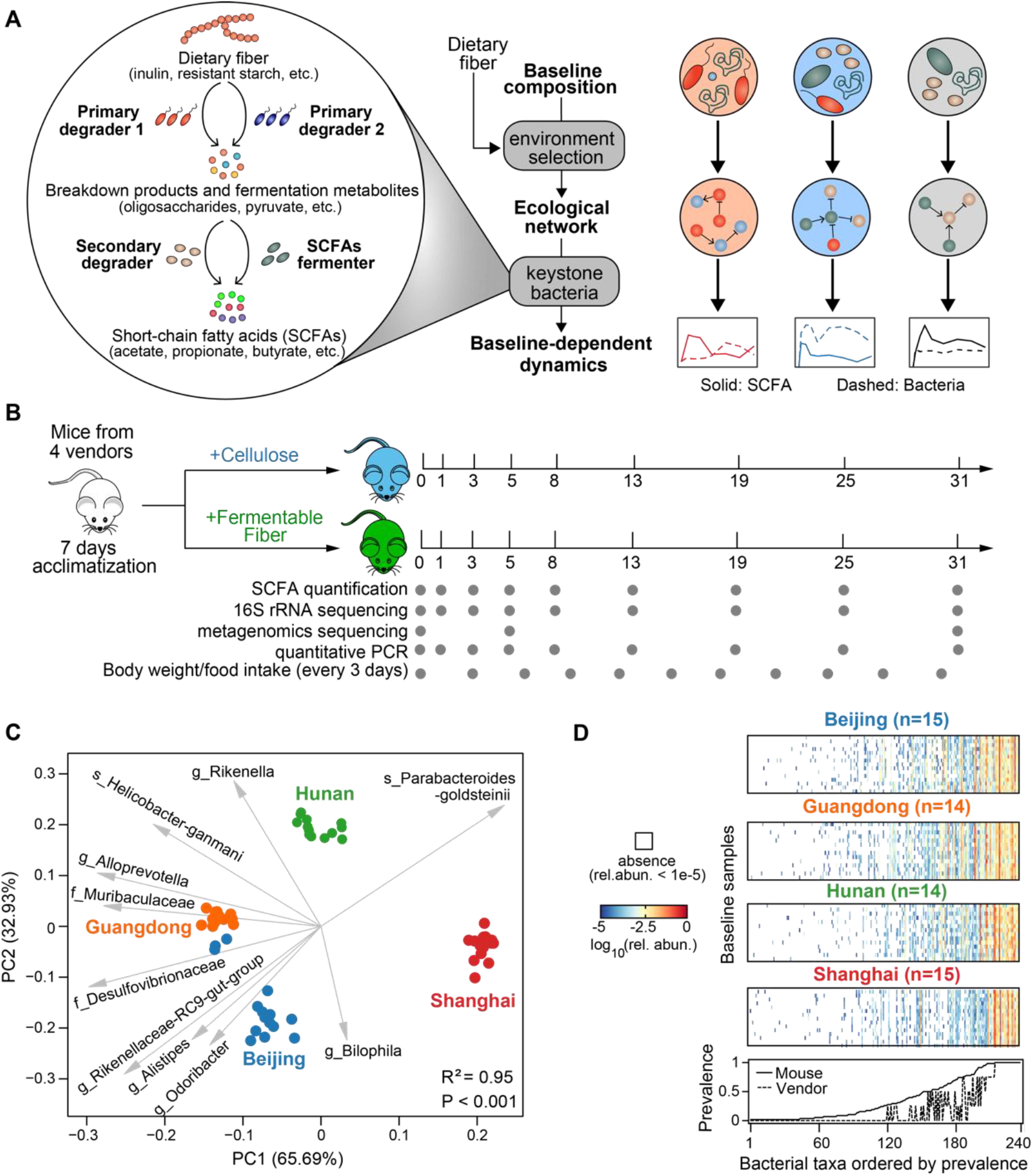
Longitudinal profiling of murine gut microbiota and metabolites to study the baselin-dependent dynamics in response to dietary fiber intervention. **A**. An ecological perspective of baseline-dependent dynamical responses of gut microbiota and SCFAs. Administration of dietary fibers alters ecological substrate niche in the gut and selects for a unique ecological network for each baseline microbiota type. Within the network, a few gut bacteria playing key metabolic roles as primary/secondary degraders and SCFAs fermenters drives heterogenous responses of bacteria and SCFAs via diverse ecological interactions (e.g., resource compeitition and cross-feeding). **B**. Experimental design. All mice from the four vendors were continuously fed with either dietary fiber (inulin or resistant starch)- or cellulose-supplemented diets for four weeks. Gray dots indicate the days on which data were collected from fecal samples. **C**. Baseline microbiota composition shown in robust PCoA (principal coordinate analysis) biplot. Isogenic age- and gender-matced mice were purchased from four different vendors (Beijing, Guangdong, Hunan, Shanghai). Gray arrows represent the dominant bacterial taxa in these samples. Adonis analysis was performed to test for differences in baseline gut microbiota composition across the four vendors (P<0.001). **D**. Top four panels: presence (white blocks indicate absence) and abundance (colored blocks) of bacterial taxa in the baseline samples. Bottom panel: the prevalence score of a taxon across mice (defined as the fraction of all mice that contains this taxon) or vendors (defined as the fraction of vendors whose mice all contain this taxon).

In this study, we profile longitudinally the gut microbiota of mice to study the ecological basis for the baseline-dependent dynamical response to dietary fibers. We use the time series data to infer the ecological network that explains why the microbiota fiber responses vary with their baseline composition. We find that the growth rates and ecological interactions of key responders—bacteria with known ability to benefit from fibers—drive the major shifts in the gut microbiota composition. And we identify the set of putative fiber-degrading bacteria whose baseline abundances predict individual responses in bacterial absolute abundance and SCFA. This study provides a framework to identify the ecological drivers of microbiota response to dietary interventions, which is critical for understanding the individualized responses of gut microbiota and the design of targeted modulations.

## Results

### Isogenic mice from different origins vary in their baseline gut microbiota

We used age- and gender-matched isogenic mice that harbor distinct baseline gut microbiota composition to study the dynamical response to dietary interventions and the inter-individual variation in ecological dynamics [28]. To ensure the distinctness of their baseline gut microbiomes, mice were purchased from four commercial vendors (labeled as Beijing, Guangdong, Hunan, Shanghai, see **Methods**), i.e. independent breeder sources. All mice were fed with cellulose-based diet 7 days prior to dietary fiber intervention. We monitored temporal shifts in the absolute abundance (by quantitative PCR) and community composition of gut bacteria (by 16S rRNA amplicon sequencing and shotgun metagenomics sequencing), SCFAs concentration (by targeted metabolomics), as well as physiological changes following the intervention of fermentable dietary fibers and cellulose (control group) (**Fig. 1B**). The two dietary fibers used in this study, inulin and resistant starch from maize, are able to be degraded by gut bacteria in the cecum/colon [29, 30] and to stimulate the production of SCFAs [8, 12].

Consistent with previous studies [31, 32], these mice grown in vendor-specific housing and feeding conditions can be naturally divided by vendor sources into groups with distinct microbiota composition after 7 days of acclimatization. Principal coordinate analysis based on the robust Aitchison distance [33] shows that the baseline compositions of those mice can be naturally clustered by vendors (Adonis, *P* < 0.001) and are characterized by distinct bacterial taxa (**Fig. 1C, Fig. S1A**). For example, Shanghai mice have low relative abundances of several commensal polysaccharide-degrading bacteria such as *Muribaculaceae* and *Rikenellaceae* [34, 35]. The profound inter-vendor differences are also noticeable at the level of presence and absence of bacteria: ∼65% taxa were entirely absent in at least one vendor and only ∼10% bacterial taxa were present in all mice (and thus all vendors) (**Fig. 1D**). Interestingly, the prevalence and abundance of bacterial taxa in these baseline samples exihibit a strong positive linear relationship in logarithmic scale (**Fig. S1B**). Due to the high between-vendor variation, mice from the same vendor can be effectively considered as independent biological replicates for each baseline microbiota composition. Throughout the observation of our experiment, the body weight of mice gradually increased over time, but the gain in body weight is generally insignificant between the inulin treatment group and the cellulose control group (**Fig. S2A**). Among the different experimental groups, there were no obvious difference in food intake and fecal weight (**Fig. S2**).

### Baseline-dependent microbiota dynamics in response to inulin

Inulin intervention rapidly promoted the absolute abundance of gut bacteria on the time scale of days, except for Shanghai mice (**Fig. 2A**). More interestingly, inulin induced a two-phase dynamical response in the gut microbiota diversity (**Fig. 2B**), which dropped rapidly in the short-term and recovered gradually in the long-term. The initial loss of diversity is primarily due to the changes in evenness (**Fig. S3A**), not richness (**Fig. S3B**), suggesting an expansion of certain bacterial taxa. Indeed, we observed rapid but non-monotonic changes in the relative abundances of several dominant bacterial genera, such as *Bacteroides* and *Muribaculaceae* (**Fig. 2C**). Notably, the long-term recovery of microbiota diversity is only partial for Beijing and Hunan mice (i.e. lower gut microbiota diversity at day 31 compared to day 0). By metagenomic sequencing, we observed temporal changes in the functional capacity of gut microbiome. Specificially, the initial (day 0), short-term (day 5) and long-term (day 31) microbiomes have distinct gene family profiles (**Fig. S4A**) and the relative abundance of genes encoding enzymes for inulin utilization (inulinases/fructanases) significantly increased after intervention (**Fig. S4B**). Collectively, our longitudinal profilings are consistent with previous observations that dietary fibers have profound impacts on the ecology and function of gut microbiota [36, 37]. In addition, we found the tendency of gut microbiota to stabilize under sustained stimulation of inulin (**Fig. 2D**). The microbiota compositions at the end of inulin intervention (all mice sacrificed on day 31) were clearly distinct from their baselines, indicating new equilibria sustained by inulin intake.

**Figure 2.**
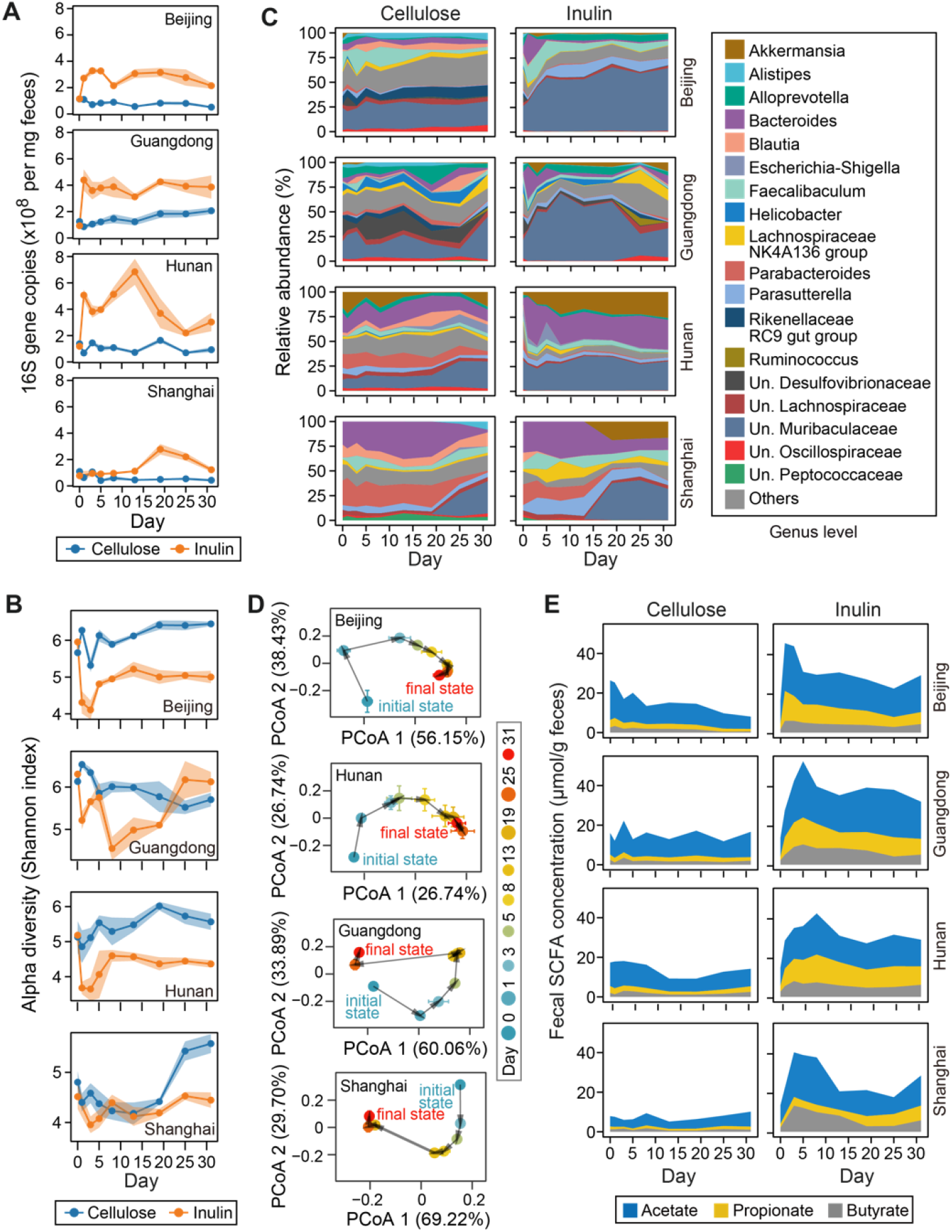
Inulin-induced temporal shifts in murine gut microbiome and short-chain fatty acids (SCFAs) metabolism. **A**. Bacterial load. **B**. Alpha diversity of gut microbiota composition. **C**. Relative abundance of bacterial genera shown in stacked band plot. Un.: unclassified. **D**. Shifts in gut microbiota composition represented by PCoA (principal coordinate analysis) plot. Initial and final states represent the microbiota compositions at day 0 and day 31 respectively. **E**. Fecal concentration of acetate, proprionate and butyrate. Beijing, Guangdong, Hunan, Shanghai are four different mice vendors. Lines (panels A,B), dots (panels A,B,D), and stacked bands (panels C, E) represent the mean values over mouse replicates from the same vendor (n=4 for Hunan and Guangdong, n=5 for Beijing and Shanghai). Shading areas (panels A,B) and error bars (panel D) represent standard error of the mean.

The changes in the gut microbiota were accompanied by changes in the levels of three major SCFAs (acetate, propionate and butyrate) and valerate (**Fig. 2E, Fig. S5**). Since SCFAs are metabolites produced by colonic bacterial fermentation of inulin, we expect a similar phase-dependent dynamics of fecal SCFAs concentration. Indeed, both total (acetate, propionate, butyrate, iso-butyrate, valerate, iso-valerate) and the three major SCFAs show two temproal phases: their levels peaked in short-term before gradually decreasing until steady states, with an exception of Shanghai mice whose propionate production was notably delayed and compromised. The mean peak-to-baseline concentration ratios of total SCFAs were 3.3, 3.9, 4.5 and 4.2 for Beijing, Guangdong, Hunan and Shanghai mice respectively. The long-term decline in SCFAs was not a result of reduced diet intake, as the intake rate remained unchanged over time (**Fig. S2**). Despite reduced SCFAs in the second phase, the mean concentrations of total SCFAs at day 31 were still 2.0-3.5 fold of its baseline levels.

We have shown above that Shanghai mice had a delayed increase in bacterial absolute abundance (**Fig. 2A**) and produced low levels of propionate (**Fig. 2E**) in response to inulin. The distinct behavior of Shanghai mice indicated that the responses of bacterial absolute abundance and SCFA production may depend on the baseline microbiota. To formally test this, we separately tested for the statistical significance of two orthogonal properties—”baseline dependence” and “responsiveness”—based on the time series data of intervention group and control group (see **Methods**). Time series data of both groups were projected onto a 2-dimensional space by sequential non-negative matrix factorization [38] to capture representative temporal trends (**Fig. 3A, Fig. S6**). With this coarse-grained data representation, we then obtained two *P*-values by comparing the differential responses between the intervention and control group (“responsiveness”, *Pr*) as well as those between the four vendors in the intervention group (“baseline dependence”, *Pb*) using Permutational Multivariate Analaysis of Variance test. This approach confirmed that the dynamical responses of bacterial load (**Fig. 3B**), propionate and butyrate (**Fig. 3C**) were nontrivial and baseline-dependent (both *Pr* and *Pb* < 0.05 after multitest correction).

**Figure 3.**
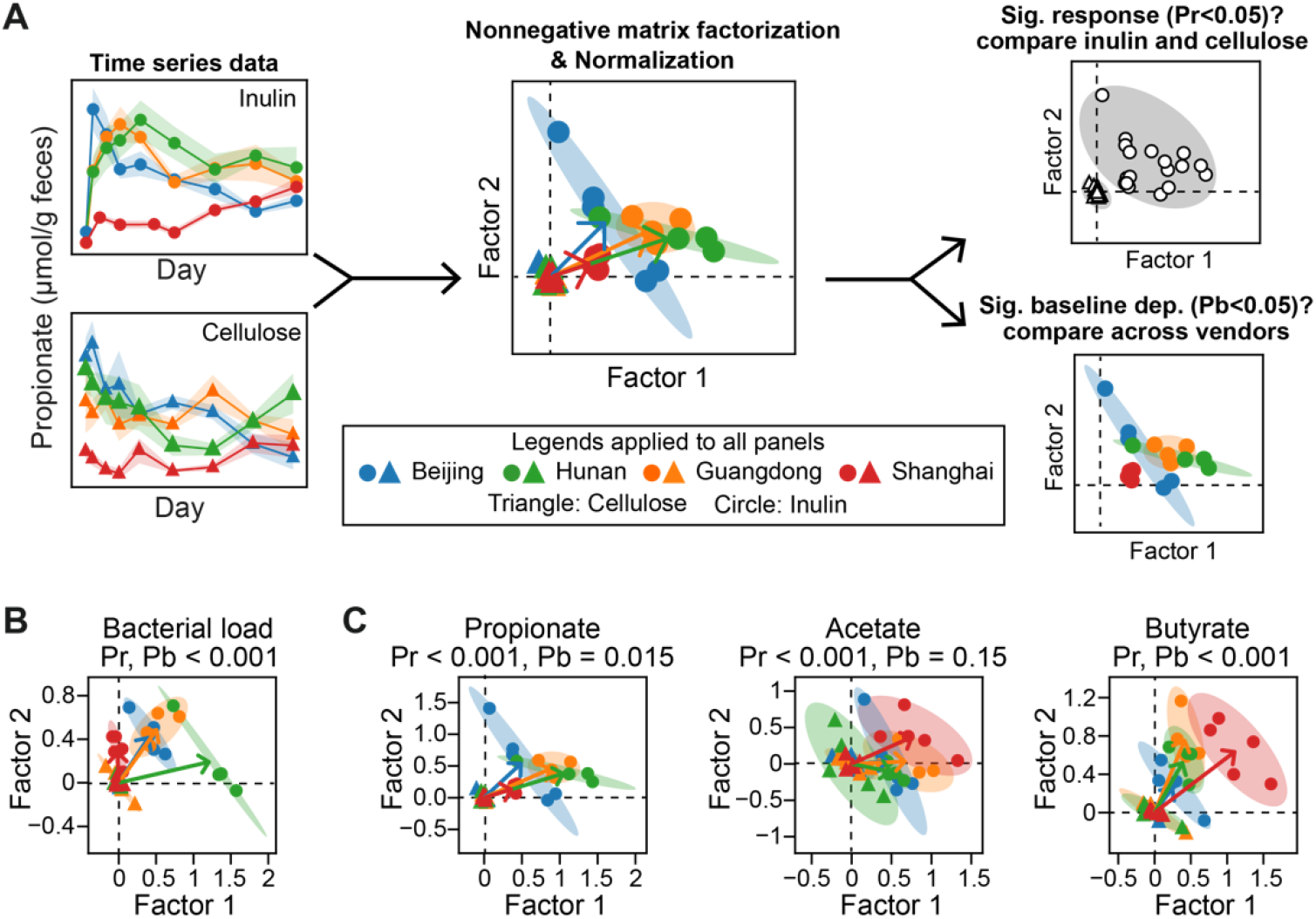
Quantifying the statistical significance of baseline-dependent dynamical response. **A**. A schematic diagram of our statistical framework to test for the significance of baseline-dependent response. The framework involves three steps: (1) projecting all time series from both intervention and control groups onto the same 2-dimensional space, (2) normalizing all data points by recentering the mean of control group to the origin, and (3) performing two separate statistical tests using the projected data to quantify the significance of “responsiveness” (*Pr*) and “baseline-dependence” (*Pb*) using Permutational Multivariate Analaysis of Variance test. Abbreviations: Significant (Sig.); dependence (dep.). For each baseline (Beijing, Guangdong, Hunan, Shanghai), an arrow was drawn from the eclipse center of the baseline under cellulose intervention (standardized to the origin) to that under the inulin intervention. **B**,**C**. Reduced 2-dimensional representation of the inulin-induced responses in bacterial load (B) and three major SCFAs (C). In all panels, each symbol represents a mouse (triangles: cellulose group, circles: inulin group) and all mice data from the same vendor under the same intervention (inulin or cellulose) was used to fit an eclipse (ellipse’s radius was determined by 2 standard deviations).

### Primary degraders of inulin respond fast and drive community dynamics

We used the generalized Lotka–Volterra (gLV) model to infer key ecological drivers of the mouse gut microbiota in response to dietary fiber intervention (**Fig. 4A**, see **Methods**). The gLV model assumes that degradation and subsequent utilization of dietary fibers boost bacterial growth rate (the amount of increment is parameterized by *ε*). One long-standing challenge in gLV inference is overfitting of parameters when the time series is too short and the number of samples at all timepoints is far less than that of gLV parameters. To address the challenge, we estimated the uncertainty associated with model parameters by formulating the gLV-based inference in a Bayesian framework which outputs posterior distributions of estimated parameters [39], rather than point estimates in penalized regressions [40]. In our gLV-based probabilistic framework, any bacteria taxa with a significantly positive distribution of “fiber response” *ε* is considered a putative “primary degrader” of inulin. We identified five such bacterial taxa grouped at varying taxonomic levels, including *Muribaculaceae* (family), *Faecalibaculum* (genus), *Parasutterella* (genus), and *Bacteroides* (genus) and *Bacteroides acidifaciens* (species) (**Fig. 4B**), see **Methods**). The inference of primary degraders in the gLV-based framework was robust to the criteria used for clustering 16S amplicons, either at the lowest classified taxonomic level (**Fig. 4B**) or at the OTU (Operational Taxonomic Unit, 97% sequence similarity) level (**Fig. S7**).

**Figure 4.**
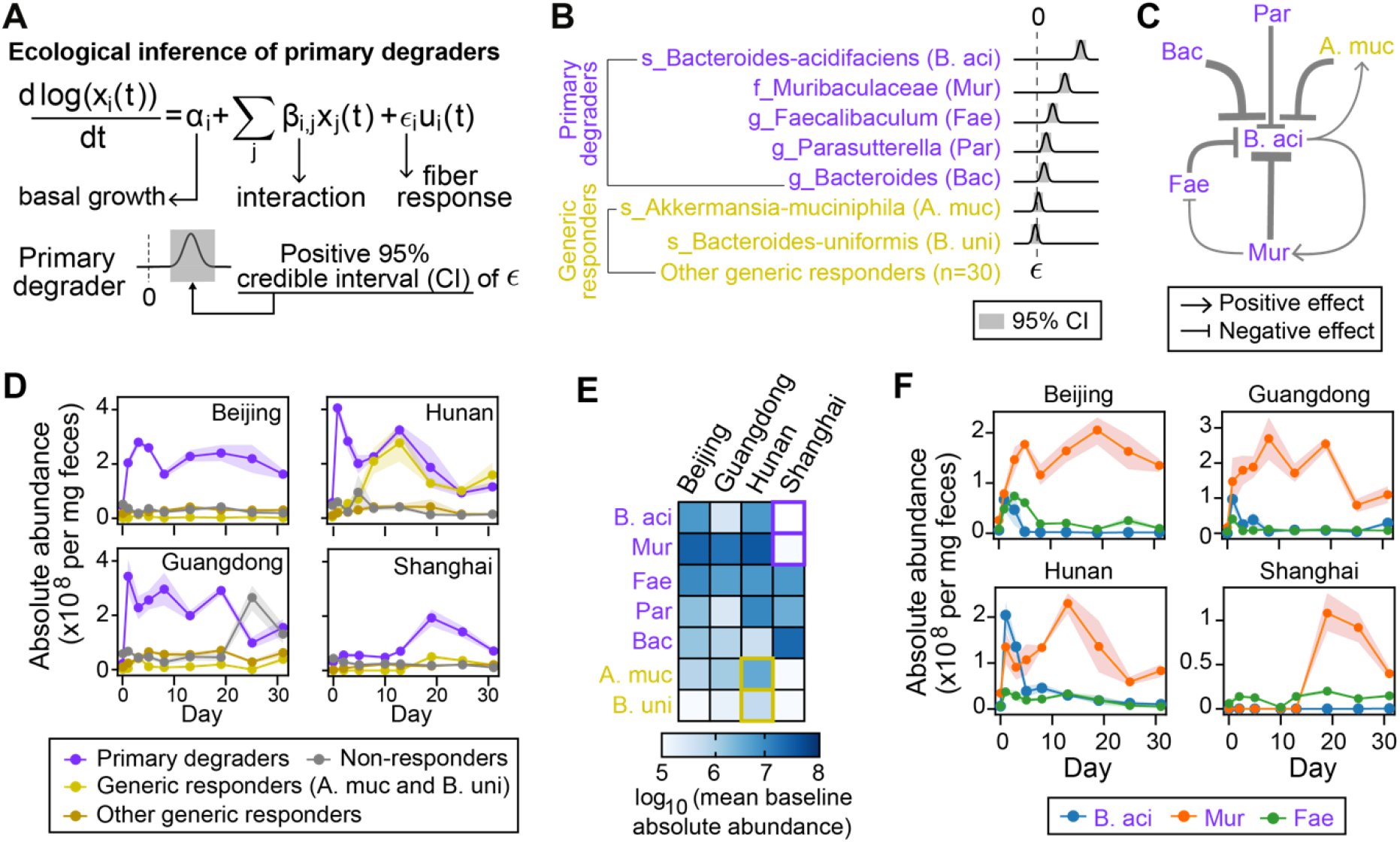
Inulin responders shape the dynamical response of murine gut microbiota. **A**. Generalized Lotka-Volterra (gLV) model combined with Bayesian statistics to infer inulin degraders and pairwise interactions. The gLV model summarizes the underlying ecology by three terms that additively determine bacterial growth rates: the basal growth rates (*α*), the influences from other bacteria (*β*), and the impacts of dietary fiber (*ε*). A primary degrader is determined when 95% credible interval of the posterior distribution of *∈* is positive. **B**. Posterior distribution of *ε* for five primary degraders (violet) and two generic responders (dark yellow). Generic responders are those bacteria showing statistical significance of inulin-induced response (i.e., responders) but not inferred as primary degraders. Bacterial taxa are ranked according to their posterior mean of *ε*. **C**. Core ecological interaction network composed of six bacterial taxa shown in the panel B (see all significant interactions in **Table S4**). Point and blunt arrows represent positive and negative interactions respectively. The arrow thickness is proportional to the posterior mean of the corresponding interaction coefficient. **D**. Ecological group dynamics of primary inulin degraders, generic responders (presented with two subgroups) and non-responders. **E**. Mean absolute baseline abundance of the seven bacterial taxa shown in the panel B. **F**. Temporal changes in the absolute abundance of the top three inulin degraders. In panels D, F, lines and dots represent the mean values across mice from the same vendor (n=4 for Hunan and Guangdong, n=5 for Beijing and Shanghai). Shading areas represent the standard error of the mean.

For four of the five putative inulin degraders (except *Parasutterella*), we found genetic evidence and/or *in vitro* experiments from literature to support their functional roles in inulin degradation (**Table S1**). For example, members from *Bacteroides* and *Muribaculaceae* contain PULs with a *susC/susD* homologous gene pair that facilitates sensing and import of inulin [41, 42]. Putative inulin PULs were also detected in the metagenome-assembled genomes of *B. acidifaciens* and *Muribaculaceae* (**Table S2**). Furthremore, we analyzed the data from an independent study [43], which profiled the murine gut microbiota composition after inulin intervention for two weeks. Analysis of this independent experiment found similar dynamics in gut microbiota diversity and composition (**Fig. S8A**,**B**). Although bacterial absolute abudance was not available in this data set, we applied the Bayesian version of gLV-based inference to the relative abundance profiles and again identified *B. acidifaciens* as a primary degrader of inulin (**Fig. S8C**).

Alternatively, we used the statistical test of responsiveness (**Fig. 3A;** *Pr*<0.05 after multitest correction) to identify all bacterial taxa that exihibited differential responses between inulin and cellulose groups, regardless of the ecological mechanism. We found a total of 37 bacterial taxa with significant dynamical responses (**Table S3**), including the five putative primary degraders as inferred by the gLV model. Among the remaining 32 “generic responders”, *Akkermancia muciniphila* and *Bacteroides uniformis* are the most abundant taxa whose relative abundances significantly increased after day 5 (**Fig. S9, Fig. S10**). Unlike the gLV-based approach, the statistical analysis on time series data did not control for indirect effects on bacterial growth via ecological interactions, so the two “generic responders” may indirectly benefit from the primary degraders. Indeed, inference of the gLV model did not identify *A. muciniphila* as a “primary degrader”, but suggested that its growth was facilitated by *B. acidifaciens* (**Fig. 4C**). This is also consistent with previous observations that *A. muciniphila* cannot grow on inulin but can be significantly promoted by shorter chain fructo-oligosaccharides [44].

The notion of eco-group (or guild), i.e., a set of bacterial taxa that perform similar functions, is very useful to understand microbial ecology [45, 46]. We divided the entire gut microbiota into three eco-groups: (1) 5 primary degraders of inulin; (2) 32 generic responders to inulin intervention; (3) non-responders. The group-level dynamics showed that the primary degraders clearly dominated the microbiota response (**Fig. 4D**). The short-term rise in the absolute abundance of a few bacterial taxa corresponds to the initial decline of the gut microbiota evenness soon after the intervention (**Fig. 2B**). More interestingly, the baseline-dependent responses can be causally linked to the initial composition of key bacterial taxa (**Fig. 4D,E**). For example, the abundances of *A. municiphila* and *B. uniformis* increased dramatically in Hunan mice (**Fig. S10**), which contained the highest abundance of these two species in the baseline (dark yellow box frames in **Fig. 4E**). Similarly, the extremely low baseline abundances of *B. acidifaciens* and *Muribaculaceae* in Shanghai mice (violet box frames in **Fig. 4E**) may explain the sluggish responses in bacterial absolute abundance and SCFA productions (**Fig. S5, S10**).

Our gLV-based inference suggested strong competition among primary degraders, where *Muribaculaceae* inhibited the growth of *B. acidifaciens* and *Facaelibaculum* (**Fig. 4C**). Indeed, *B. acidifaciens* and *Facelibaculum* showed transient dynamics with a quick rise and drop in their absolute abundances, while the abundance of *Muribaculaceae* increased steadily and remained high during the entire period of observations (**Fig. 4F, S10**). Our results are consistent with previous studies by Patnode *et al*. [47] that identified competitive inhibition as the ecological mechanism for consistent dominance of *Bacteroides cellulosilyticus* over *Bacteroids vulgatus*, even though both species contain fiber-processing polysaccharide utilization loci (PULs). Taken together, we demonstrate that primary degraders and their competitions are key drivers of the baseline-dependent ecological dynamics of microbiota response to dietary fibers.

### Baseline-dependent SCFA production and its association with gut microbiota composition

The dynamics of SCFAs during inulin intervention varied substantially across different baselines (**Fig. S5**). Shanghai mice produced the lowest level of propionate (**Fig. S5**); these mice also showed the lowest response in bacterial load (**Fig. 2A, Fig.4D**), due to very low abundance of some primary degraders and generic responders of inulin in the baseline microbiota (**Fig. 4E**). We hypothesized that these key taxa may directly contribute to propionate production and found that the baseline abundances of *B. acidifaciens, Muribaculaceae, A. municiphila, B. uniformis* were positively correlated the propionate concentration (**Fig. 5A**, left panel). Indeed, *Muribaculaceae, A. municiphila* and *B. uniformis* have been previously found to produce propionate *in vitro* and/or *in vivo* (**Table S1**). As a result and consistent with a previous study [48], there was a strong positive association between bacterial load and propionate concentration (**Fig. 5A**, right panel; P<0.001), as the two are both baseline-dependent. In contrast, the association between bacterial load and other SCFAs was not significant (**Fig. S11**).

**Figure 5.**
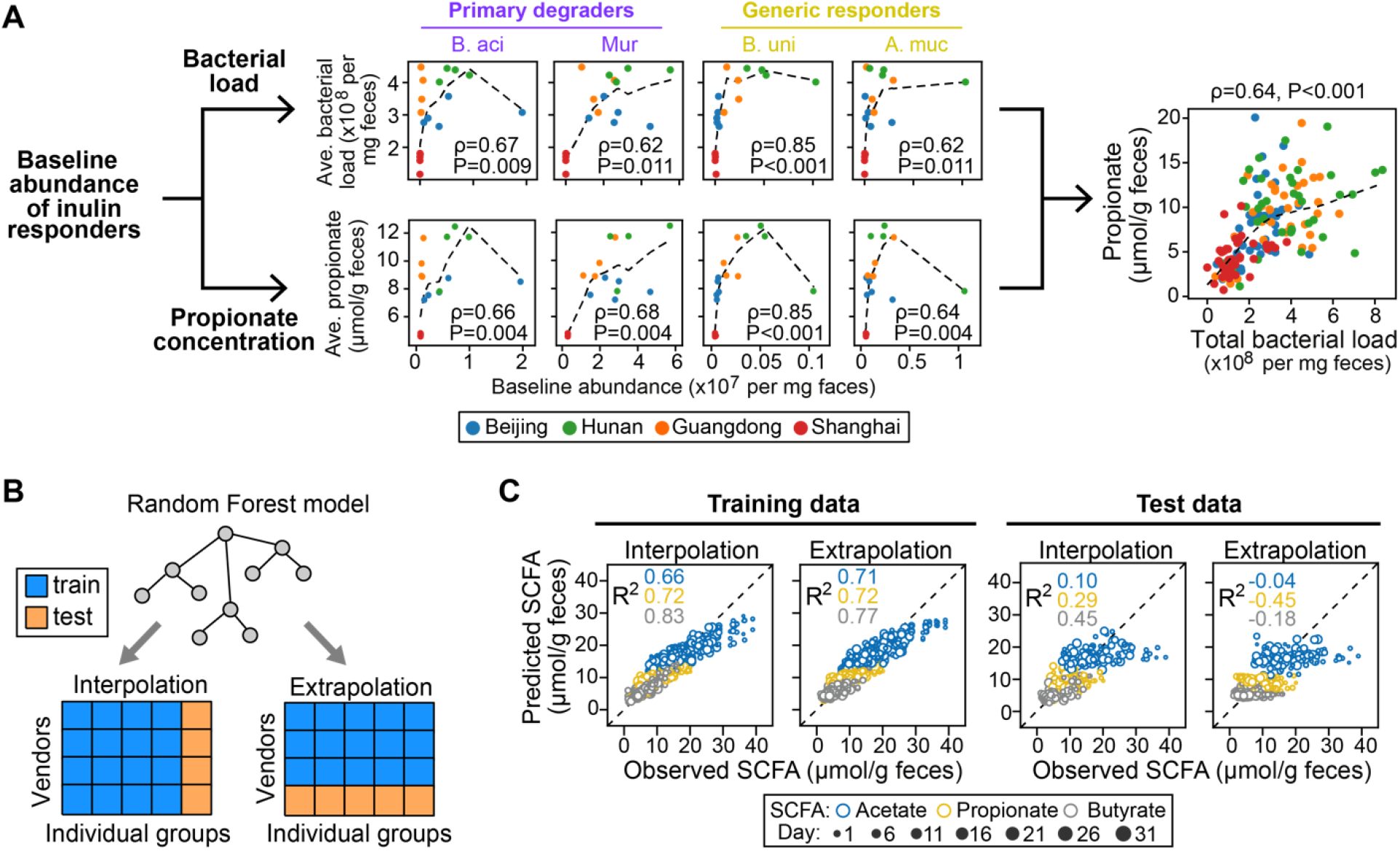
Quantitative relationship between SCFAs and murine gut microbiota composition. **A**. Correlation between bacterial load and propionate concentration (right big panel). We proposed that the correlation is mediated by some inulin responders which causally and simultaenoulsy affect both observations. Eight small panels to the left: Spearman’s correlations of baseline abundance of four inulin responders (B. aci: s_Bacteroides-acidifaciens, Mur: f_Muribaculaceae, B. uni: s_Bacteroides-uniformis, A. muc: s_Akkermansia-muciniphila) with the mean bacterial load (top row) or propionate concentration (bottome row) averaged across the interveion period. Dashed line: Lowess (Locally Weighted Scatterplot Smoothing) regression. Spearman correlation coefficient (ρ) and adjusted P-value are indicated in each plot. **B**. Prediction of SCFAs concentrations from gut microbiota composition using machine learning models. Two data-split strategies for testing model performance were designed: mice in the test sets were randosmly selected on a one-per-vendor basis for “interpolation” and exclusively selected from a single vendor for “extrapolation”. Data before intervention (i.e., day 0) was not included. **C**. Performances of Random Forest regression models on the training and test datasets.

Given that the gut microbiota is strongly associated with the fecal levels of fiber fermentation products, we asked whether we could quantitatively predict SCFA concentrations from the microbiota composition measured at the same time. We evaluated the performance of machine learning models to predict the fecal SCFA concentrations using absolute abundance of bacterial taxa as predictors. All mice in our experiments were split into training data and test data using different data-split approach (**Fig. 5B**). The “interpolation” approach generated balanced distribution of baseline microbiota composition between the training and test data (**Fig. S12A**), by randomly selecting a single mouse from each vendor as test data and using the other mice for training. In contrast, the “extrapolation” approach produced highly unbalanced microbiota distribution between the training and test data (**Fig. S12B**), by randomly selecting all mice from a vendor as test data and using mice of the other vendors for training. Although the Random Forest regression model fitted the training data reasonably well (R^2^ ≥0.66 regardless of SCFAs and data-split approaches), the predictions generalized poorly to the test data: R^2^ of SCFAs ranged from 0.1 to 0.45 for “interpolation” but dropped below 0 for “extrapolation” (**Fig. 5C**). We further showed that the low predictability in extrapolation cannot be substantially improved by using alternative predictors (**Fig. S13A**), models (**Fig. S13B**) or adding weights to training samples (**Fig. S13C**). Given the current sample size, we found that Random Forest regression models based on gut microbiota composition had low or no predictive power for fecal SCFA concentration, if the gut microbiota of interest was significantly different from the baselines covered in training data. This agrees with previous studies in humans finding[49] that the substantial inter-individual variation of gut microbiome could impede the predictive power of machine learning models.

### Response to resistant starch validates model developed from response to innulin

To study whether our ecological framework can be generalized to study the dynamical responses of gut microbiota to other dietary fibers, we administered resistant starch from maize to mice from the same four vendors following the same experimental procedure (see **Methods, Fig. 1B**). Compared to inulin, resistant starch stimulated milder changes in the bacterial load (**Fig. 6A**), gut microbiota composition (**Fig. 6B**), and SCFAs production (**Fig. 6C**). Under resistant starch intervention, we identified *Faecalibaculum* and *Muribaculaceae* as “primary degraders” (**Fig. 6D**, see **Table S1** for genetic evidence from literature) and 25 additional bacterial taxa as “generic responders” (**Table S3**). The dynamics of primary degraders were qualitatively similar between inulin and resistant starch interventions (**Fig. 6D**): *Muribaculaceae* increased rapidly and reached a plateau (except for Shanghai mice), while *Faecalibaculum* declined sharply after the initial burst. The gLV-based inference suggested that the observed dynamics was driven by mutual inhibition between the two primary degraders (**Fig. 6E**).

**Figure 6.**
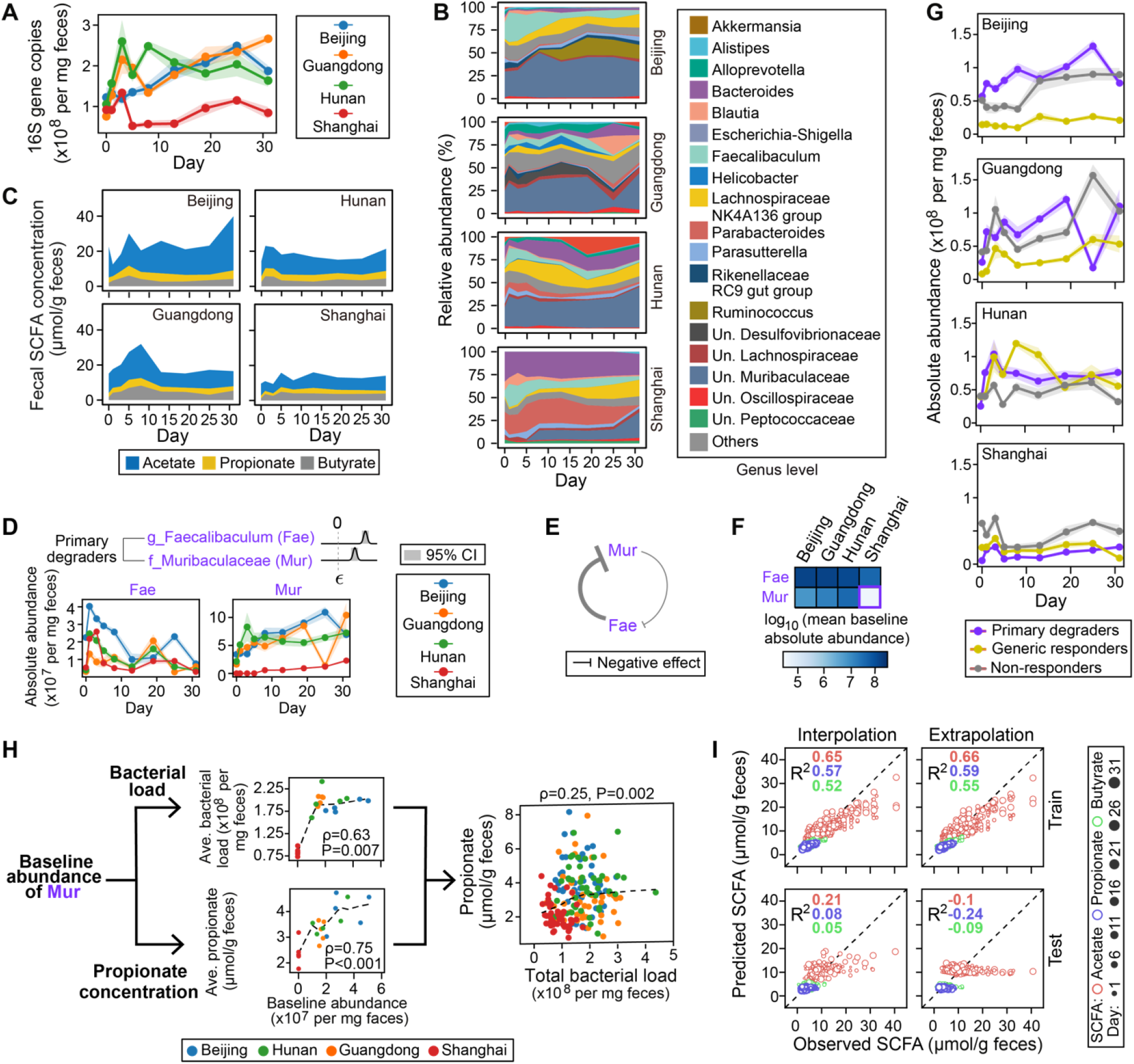
Resistant starch-induced dynamical response in murine gut microbiota. **A-C**. Dynamical responses of bacterial load (A), gut microbiota composition (B), and SCFAs concentration (C) following resistant starch intervention. **D**. Dynamics of two putative resistant starch degraders. *∈* represents the growth impact of resistant starch and its posterior distributions are shown for each degrader. CI: credible interval. **E**. Ecological interactions between the two degraders (see all significant interactions in **Table S4**). The arrow thickness is proportional to the posterior mean of the corresponding interaction coefficient. **F**. Mean baseline abundances of the two degraders. **G**. Ecological group dynamics of primary degraders and generic responders of resistant starch as well as the non-responders. **H**. Correlations among baseline abundance of *Muribaculaceae*, bacterial load and propionate concentration. For the two small panels on the left, the time-averaged bacterial load or propionate concentration on the y-axis were calculated by dividing the area under the curve of the corresponding variables by the duration of observation. Dashed lines: Lowess (Locally Weighted Scatterplot Smoothing) regression. **I**. Prediction of SCFAs concentration from gut microbiota compsotion using a Random Forest regression model. “Interpolation” and “extrapolation” are two strategies of splitting all data into the training and test sets (the same as **Fig. 5B**). Lines (panels A,D,G) or height of stacked bands represent mean values across mice from the same vendor (n=5 for all vendors). Shading areas (panels A,D,G) represent the standard error of the mean.

We found that bacterial load (**Fig. S14A**) and the three major SCFAs (**Fig. S14B**) exihibited baseline-dependent dynamical responses to resistant starch intervention. For example, the weak response in bacterial load and SCFA production of Shanghai mice (**Fig. 6A**) could be explained by the low abundance of *Muribaculaceae* in the baseline community (**Fig. 6F**, highlighted in red box frame). In addition, there was substantial growth of generic responders in Hunan mice (**Fig. 6G**), although the dominant bacterial taxa in this eco-group were different from the taxa identified in inulin intervention (**Table S3**).

Finally, we found that the baseline abundance of *Muribaculaceae* was separately correlated with bacterial absolute abundance and prioprionate level (**Fig. 6H**, left panel), supporting the hypothesis that *Muribaculaceae* may serve as both a primary degrader and a propionate producer. As a result, there was a weak but statistically significant positive association between bacterial load and propionate concentration (**Fig. 6H**, right panel; P=0.002). Similar to our findings from the inulin intervention group, Random Forest models based on gut microbiota compostion had low or no predictive power for SCFA concentration in the resistant starch intervention group (**Fig. 6I**). Collectively, our major findings were qualitatively consistent between inulin and resistant starch interventions, suggesting that the dynamical responses of gut microbiota to fiber-based perturbation follow universal ecological principles.

## Discussion

Our study characterizes the ecosystem response of murine gut microbiota to fibers, and emphasizes that ecological interactions play a key role in the personalized impact of dietary changes. gLV-based ecological inference from gut microbiome time series data has yielded mechanistic insights into the stability of probiotic community under dietary perturbation [49], colonization resistance of pathogenic *Clostridioides difficile* [51], and community assembly dynamics within preterm infant gut [24]. By integrating gLV model with Bayesian regression, we inferred a competitive network of fiber degraders as key bacteria that mediate the response of murine gut microbiota to inulin and resistant starch intervention. Besides evidences supporting the fiber-degrading function of putative degraders, our study confirms findings in the literature and advances the understanding of the effects of dietary fibers on the gut microbiome at the system level. First, the small number of fiber degraders (five for inulin and two for resistant starch) suggested that fiber-induced bacterial shifts are very selective and occur to a restricted number of taxa. Second, the absolute abundance of many fiber-degrading bacteria such as taxa related to the genus *Bifidobacterium*, failed to expand in the mouse gut on both fibers (**Fig. S15**), indicating that fiber-induced bacterial enrichment cannot be simply predicted from their *in vitro* growth. Third, we reasoned that the personalized fiber-induced response of gut microbiota were largely determined by the baseline abundance of fiber degraders and the ecological interactions among these degraders. In our study, the inter-individual variability of fiber-induced shift in propionate production was associated with the baseline abundance of a key fiber degrader *Muribaculaceae*. This is consistent with previous findings that the abundance of a key starch-degrader prior to resistant starch intervention was indicative of whether an individual would have higher fecal butyrate response [16]. The ecological interactions among fiber degraders, on the other hand, drives the cascading alterations during the intervention. Our results revealed that fiber-induced dynamics of murine gut microbiota were largely driven by competitions and *Muribaculaceae* outcompeted other degraders in consuming both inulin and resistant starch. Since the family *Muribaculaceae* was specific to the mouse gut [52], it might have been adapted to the murine gut with higher fitness.

Understanding the microbial responses and its association with the baseline microbiota composition rightly is a critical step in individualized dietary fiber intervention. To date, most of the current studies are based on cross-sectional study design and described the pre-to-post changes in abundance/concentration as microbial responses to dietary intervention [13, 15]. However, it should be noted that gut microbiome is a highly dynamic ecosystem, and its response to dietary fibers could have temporal characteristics [37]. Consequently, the significance of microbial responses may vary depending on the study endpoint used to calculate pre-to-post changes. In our experiments, the changes in propionate concentration from their baseline levels differ significantly among the four vendors at day 5 but not at day 31 (**Fig. S16**). Furthermore, due to the lack of control group data to assess the intervention effects, pre-to-post changes that are supposedly to capture fiber-indcued effects may be entirely attributable to random temporal variations within each individual [53]. We speculate that these two caveats are the main cause of “reproducibility crisis” [54] among microbiome researches. In constrast, our analysis avoids the above two caveats by incorporating longtitudinal data and a control group. Additionally, the use of dimensionality reduction in our approach further benefits data visualization of inter-vendor variations in gut microbiota composition (**Fig. 3**).

Diet-induced changes in SCFAs are often transient and vanish shortly after cessation of dietary intervention [9, 55-57]. Our study is consistent with this result, by showing that SCFA concentrations cannot be maintained at its peak and drop by 35%-40% even under continuous inulin intake until four weeks. The transient responses in SCFAs were also observed in colorectal cancer patients [58] and type 2 diabetes mellitus patients [6]; however, it is unknown whether the reduced SCFAs in these human subjects are resulted from lower dietary fiber intake. Despite the drop, our data demonstrates that a continuous intervention that lasts for 31 days is sufficient to elevate and stabilize the SCFAs concentration, but it is not clear yet whether the elevated level persists after the intervention discontinues. The *in vivo* SCFAs dynamics is jointly determined by multiple metabolic processes, where the two major ones are microbial production and host absorption. In healthy individuals, 90%-95% SCFAs produced in the colonic lumen are absorbed by the gut mucosa [59]. While many studies used fecal SCFAs concentrations as proxy of their luminal levels, neither of both represents the rate of production or absorption so the declined phase of SCFAs in our study may be explained by reduced production rate, increased absorption rate or both. Due to the difficulty of measuring fluxes *in vivo*, mathematical models that take both processes into accounts show great premise in the estimation of their flux rates from SCFAs concentrations [60].

Characterizing the dynamics of gut microbiota and their inter-individual variability with multi-omics data is an important priority for microbiome research to further understanding of diet-induced responses [61]. Such studies hold great promise to improve human health and treat gut microbiome-associated disease via microbiome engineering. A key question in microbiome engineering with prebiotics is whether and to what extent can we enrich the gut levels of beneficial bacteria using prebiotic compounds. Microbiome engineering, as with other engineering disciplines, requires computational tools to aid the design process. Predicting bacterial responses to interventions in the human gut is nontrivial: previous studies have repeatedly shown that bacteria able to consume a fiber supplement *in vitro* may not be selectively enriched *in vivo*, suggesting that dietary response of a gut bacterial taxa depends on the ecological context [62]. By inventing a new application of gLV with uncertainty assessment to infer primary fiber degraders and the associated interaction network, we provide a generalizable computational approach to study the ecological dynamics of the gut microbiome under dietary interventions. We foresee that applications of ecological modeling in human cohorts with dense longitudinal sampling will provide important insights for predictable dietary responses and personalized nutrition [63].

## Methods

### Animal experiments

Specific-pathogen-free (SPF) female C57BL/6J mice were obtained at 6 weeks of age from four different vendors, including Beijing (A Charles River Company, Beijing, China), Hunan (Hunan Slac Jingda Laboratory Animal Company, Ltd., Changsha, China), Guangdong (Guangdong Medical Laboratory Animal Center, Foshan, China)) and Shanghai (SLAC Laboratory Animal Co., Ltd., Shanghai, China). Mice were maintained in 12-h light/dark cycle and allowed ad libitum access to food and water throughout the experiment. After acclimatizing to the diet and housing environment for 1 week, mice from each vendor were randomly separated into three groups: cellulose group (n = 5), resistant starch group (n = 5), and inulin group (n = 5). Composition of all diets including the source of dietary fibers cellulose, resistant starch from maize (HI-MAIZE® 260, Ingredion Inc.), and inulin (Orafti HP, BENEO-Orafti) are provided in **Table S5**. Fecal pellets from each mouse were freshly collected over multiple time points: day 0 (before diet change), day 1, 3, 5, 8, 13, 19, 25, and 31 (Figure 1A). Fecal samples were snap-frozen in liquid nitrogen and stored at −80 °C until further processing. At every cage change (moving the mice to a new clean cage with fresh bedding twice in one week), body weight was individually measured, and food intake and fecal output of each cage mice during the previous three days per cage were measured. This study was approved by the Institutional Animal Care and Use Committee of the Shenzhen Institutes of Advanced Technology, Chinese Academy of Sciences.

### Quantification of fecal SCFA concentration by GC-MS

The SCFAs of mice fecal samples were analyzed by GC-MS [63]. For the sample extraction, 0.05 g of frozen feces were mixed with 300 µL of pure water containing caproic acid-6,6,6-d3 (CDN Isotopes, Quebec, Canada) as internal standard (final concentration 20 µg/mL). After adding 1.0 mm diameter zirconia/silica beads (BioSpec, Bartlesville, OK), feces were homogenized for 20 s under 6500 rpm for three times, then incubated at 4 °C with shaking for 30 min, followed by centrifugation for 30 min at 13,000×g. Following extraction with anhydrous diethyl ether, the SCFA extract accurately transferred into a glass insert in a GC vial and capped tightly after added 5 µl of N, O-bis(trimethyl-silyl)-trifluoroacetamide and vortexed for 5 s. The mixture was kept in the GC vial and incubated at room temperature (22 °C) overnight (or over 8 h) before loading to GC/MS. The analysis of acetic, propionic, iso-butyric, iso-valeric, valeric and butyric acids was performed by Agilent 8890/7000D triple quadrupole GC/MS equipped with a capillary HP-5 ms capillary column (30 m × 0.25 mm × 0.25 µm film thickness) (Agilent Technologies). The analytes were quantified in the selected ion monitoring (SIM) mode using the target ion and confirmed by confirmative ions. The integrated areas for all SCFAs were normalized with the internal standard and quantified with the standard curve constructed.

### DNA extraction and quantification of bacterial load

DNA of mice fecal samples was extracted using the QIAmp PowerFecal DNA kit (Qiagen, #12830–50) following standard manufacturer procedures. DNA samples were resuspended in Buffer C6 and quantitated using the Qubit fluorometer (ThermoFisher Scientific). To quantitatively assess bacterial load, total bacteria density were determined using qPCR as previously described [66]. A 466-bp fragment of the bacterial 16S ribosomal DNA was amplified using the forward primer 5′-TCCTACGGGAGGCAGCAGT-3′ and the reverse primer 5′-GGACTACCAGGGTATCTAATCCTGTT-3′. The absolute abundance of a bacterial taxon was estimated by multiplication of its relative abundance and the total bacterial load.

### 16S rRNA amplicon sequencing and shotgun metagenomic sequencing

16S rRNA gene sequencing was performed as previously described with modifications [67]. Library preparation was done using a two-step PCR method. During the first step of PCR, primers S-D-Bact-0341-b-S-17 (forward) and S-D-Bact-0785-a-A-21 (reverse) were used to target and amplify the v3-4 region [68], as well as to add second-step priming sites. Dual index codes were added to each sample at the second PCR step. The PCR products were purified with Agencourt AMPure XP magnetic beads (Beckman Coulter, Brea, CA, USA) and quality controlled with TapeStation (Agilent Technologies, Santa Clara, CA, USA). The final DNA concentrations of the purified products were measured with Qubit 2.0 fluorometer (Thermo Fisher Scientific). The purified products were pooled in equal molar concentrations, and denatured following the Illumina protocol. Sequencing was performed in a single run on NovaSeq 6000 (Illumina, USA). Blank controls (no sample added, processed routinely, n = 4) were included in the extraction process to control for contamination throughout processing.

Metagenomic sequencing was performed using fecal samples from the inulin diet group at day 0, 5 and 31. Extracted DNA sample was purified using silica-based columns. Metagenomics sequencing libraries were prepared with at least 2 μg of total DNA using the Nextera XT DNA sample Prep Kit (Illumina, San Diego, USA) with an equimolar pool of libraries achieved independently based on Qubit 2.0 fluorometer results combined with SYBR Green quantification (Thermo Fisher Scientific, Massachusetts, USA). The indexed libraries were sequenced on NovaSeq 6000 (Illumina, USA) by Guangdong Magigene Biotechnology Co.,Ltd. (Guangzhou, China).

### Bioinformatics analysis

The 16S rRNA sequencing reads were analyzed by QIIME 2 (version 2020.2) [69]. Demultiplexed paired-end reads were trimmed to remove primers and low-quality bases with q2-cutadapt plugin. The trimmed sequences were denoised and joined with q2-dada2 plugin. Potential reagent contaminants were identified using decontam package based on either the frequency of the amplicon sequence variants (ASVs) in the blank control or the negative correlation with DNA concentration [70]. The generated feature table was filtered to remove ASVs present in only a single sample and remaining ASVs were used to construct a rooted phylogenetic tree via q2-phylogeny. Rarefaction curve analysis of the data obtained was used to estimate the completeness of microbial communities sampling and performed using the iNEXT R package [71]. Subsequently, in order to avoid sample-to-sample bias due to variable sequencing depth (different number of reads per sample), samples were rarefied to 38,980 sequences per sample. Rarefaction analysis showed that great majority of the bacteria species diversity and richness that could be sampled was captured by our sequencing depth (**Fig. S17**), indicated sufficient sequencing depth for majority of the analyzed samples. Estimated alpha diversity metrics using q2-diversity. Beta diversity metrics (Aitchison distance) and biplot were generated using DEICODE (robust Aitchison PCA, RPCA) [33]. Group significance between alpha and beta diversity indexes was calculated with QIIME2 plugins using the Kruskal–Wallis test and permutational multivariate analysis of variance (PERMANOVA), respectively. To assign taxonomy to ASVs, the q2-feature-classifier basing on the classify-sklearn naïve Bayes taxonomy classifier was used with the SILVA (v.138) as reference database. Unless specified (**Fig. 2C, Fig.6B, Fig. S1A, Fig. S7**), ASVs were grouped to the lowest classified taxonomy level (i.e., grouping 16S sequencing at a specified taxonomic level, excluding those classified at lower levels) for all data modeling and analysis. The specified taxonomic levels are labelled with prefix “s_” (species level), “g_” (genus level), “f_” (family level), “o_” (order level), “c_” (class level), “p_” (phylum level), and “k_” (kingdom level) to indicate the taxonomic rank where grouping was operated. For example, “g_Bacteroides” clusters all sequences that are classified as Bacteroides at the genus level but unclassified at the species level. Alternatively, ASV sequences were grouped into Operational Taxonomic Units (OTUs) at 97% similarity (**Fig. S7)**.

For metagenome analysis, raw sequencing reads were subjected to quality filtering and barcode trimming using KneadData (v0.5.4) by employing trimmomatic settings of 4-base wide sliding window, with average quality per base >20 and minimum length 90 bp. Reads mapping to the mouse genome were removed. Kraken2 was run against genome taxonomy database (GTDB_r89_54k) with default parameters [72]. Following classification by Kraken2, Bracken was used to re-estimate bacterial abundances at taxonomic levels from species to phylum using a read length parameter of 150. Next, the filtered sequences were assembled into contigs using metaSPAdes with default settings [73]. The gene abundance was analyzed and calculated as previously described with modifications [74]. Putative genes were then predicted on contigs longer than 200 base pairs using Prodigal under metagenome mode (-p meta) [75]. A non-redundant gene catalogue was constructed with CD-HIT using the parameters “-c 0.95 –aS 0.9” [76]. The abundance of each predicted gene was evaluated by mapping reads back with KMA algorithm and then normalized with the following equation: RPM = 1M × (mapped reads/gene length)/(sum of mapped reads/gene length) [77]. For all the predicted genes, CAZymes were annotated using hmmsearch against the dbCAN2 database V9 (e value <1 × 10−10; coverage >0.3) [78]. The domain with the highest coverage was selected for sequences overlapping multiple CAZyme domains. For all samples, short genomic assemblies (<2,000 bp) that could have biased the subsequent analysis were first excluded. Genomes were then binned using VAMB [79]. The binning results were refined based on the bin quality assessment (completeness >75, and contamination <15) of different binners from CheckM [80]. Taxonomic classification of each bin was determined by GTDB-tk [81], and subjected to prediction of polysaccharide utilization loci (PULz) using pipeline PULpy [82].

### Significance test of baseline-dependent responses

Sequential non-negative matrix factorization [38] was applied to transform all high-dimensional time series data from both intervention (inulin and resistant starch) and control group into two-dimensional space. We chose two factors because (1) reconstructed time series from the two latent factors preserve the quantitative trends of the untransformed time series sufficiently well and (2) two-dimensional data can be easily visualized. The reduced representation of the intervention group {(*x*_*v,i*_, *y*_*v,i*_)} and control group {(*p*_*v,j*_, *q*_*v,j*_)}, *v* (*v* = 1,2, …, *V*) refers to the index of vendor and *i, j* (*i* = 1,2, …, *N* and *j* = 1,2, …, *N*) refers to the index of mouse. For each vendor *v*, both vectors were then standardized by subtracting the mean vector of the vendor in the control group, i.e., 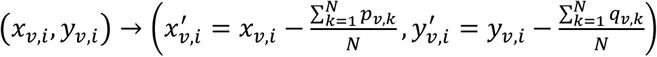 and 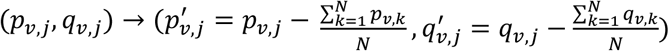.

The statistical significance test of: 1) the responsiveness (i.e., whether time series in the intervention group 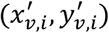 differs from that in the control group 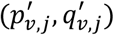 regardless of vendor), and 2) the baseline dependence (i.e., whether time series in the intervention group 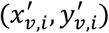 varies among vendors *v*), were performed separately using Permutational Multivariate Analaysis of Variance (PERMANOVA) with Minkowski distance as the distance metric (the statistical module “statsmodels” in Python). We then obtained two P-values by comparing the differential responses between the intervention and control group (“responsiveness”, Pr) as well as those between the four vendors in the intervention group (“baseline dependence”, Pb). If both P-valus are smaller than 0.05, we determined that a quantity has a baseline-dependent response. For all significance tests that require multiple test correction, the Benjamini-Hochberg procedure [83] was used for controlling the false discovery rate in multiple test correction.

### Ecological inference of dietary fiber responses

The generalized Lotka-Volterra (gLV) model describes how the absolute abundance of bacterial species change over time

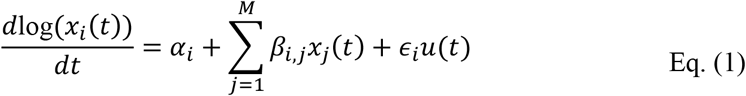

where *M* is the number of bacterial taxa, *x*_*i*_ is the absolute abundance of taxon *i* (*i* = 1,2, …, *M*), *α*_*i*_ is the basal growth rate, *β*_*i,j*_ represents the influence of taxon *j* (*j* = 1,2, …, *M*) on the growth of taxon *i, ∈*_*i*_ is the susceptibility coefficient that represents growth response to dietary fiber, *u*(*t*) is a binary variable that indicates whether the fiber is administed at time *t*. Bayesian regression techniques were used to parameterize the generalized Lotka-Volterra (gLV) model, as similarly used in Morjaria et al [39]. For each mice *r* (*r* = 1,2, …, *P*), Eq. (1) can be transformed into a matrix form that incorporates all discrete time points of measurements (*t*_*k*_, *k* = 1,2, …, *N*)

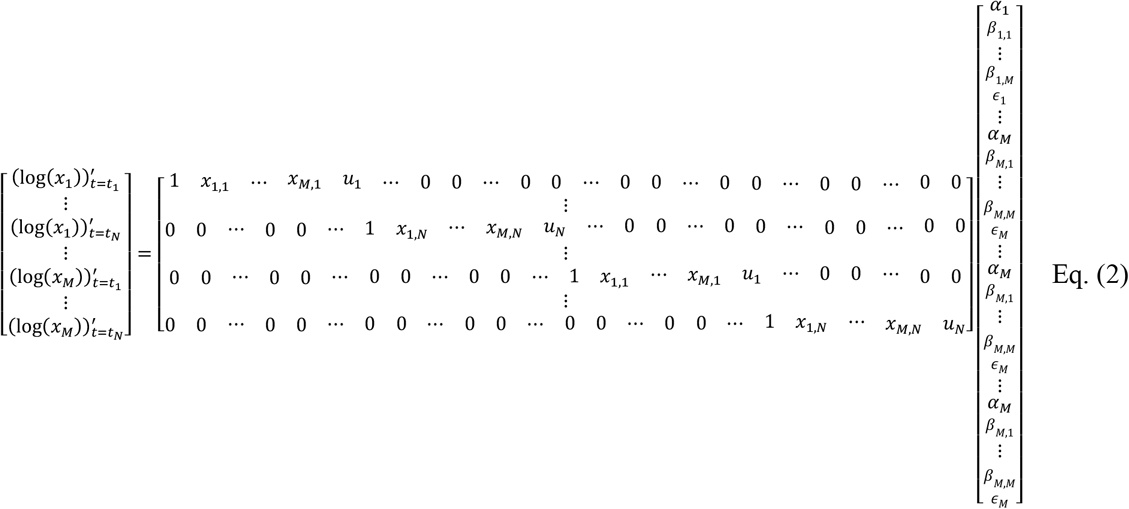

where *x*_*i,k*_ = *x*_*i*_(*t*_*k*_) and *u*_*k*_ = *u*(*t*_*k*_). The log-derivatives of *x*_*i*_ on the left-hand side of Eq. (2) were estimated from a cubic spline interpolation. Using a simplified notation for Eq. (2), i.e., **Y**_***r***_ = **X**_***r***_**C**_***r***_, we can incorpates data from all mice into a single regression model

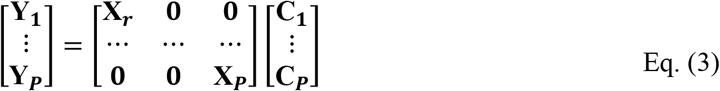

The linear regression as described in Eq. (4) (for brevity **Y** = **XC**) can be further transformed into a Bayesian regression **Y** = 𝒩(**XC**, σ) where 𝒩 and σ represent normal distribution and standard deviation respectively.

Since gLV models the absolute abundance of bacterial taxa, we multiplied the bacterial load by their relative abundance to calculate absolute abundance. The time-series data from all mice were simultaneously fed into the gLV model based on the premise that ecological forces driving microbiome dynamics are largely host-independent and universal [84]. We used uninformative Normal priors 𝒩(0,1) for all gLV parameters and Stan program [85] to produce posterior distributions for each parameter after “no U-turn” sampling of 10,000 samples from at least 3 independent Markov chain Monte Carlo traces. Since Stan is computationally expensive, we limited the inferences of dietary fiber responders to the top 20 bacterial taxa with the highest time-averaged absolute abundances in the inulin (or resistant starch) and cellulose group. Our Bayesian approach is conceptually similar to the Bayesian adaptive lasso algorithm in MDSINE [50]; the key difference is that MDSINE uses a hierarchical probability model and regularization to sample the variance parameters in the Normal priors.

### Random forest (RF) model

Model development was run in a pipeline by combining normalization for data transformation, LASSO (least absolute shrinkage and selection operator) for feature selection, and RF regression for data fitting and prediction. The tolerance used in LASSO is 1e-5 and features whose coefficients below this threshold were discarded and not used to build RF regression model. Two data-split approaches were implemented. For the “interpolation” approach, the mice from the same vendor were first alphabetically labeled as A-D (for Hunan and Guangdong) or A-E (for Beijing and Shanghai). Then the four mice with the same label (one per vendor) were chosen to constitute the test set and the remaining mice were mixed together to form the training set. For the “extrapolation” approach, all mice from a specific vendor were chosen as the test set while the training set includes all mice from the other three vendors. To train each model, five hyperparameters were tuned using 5-fold cross validation within the training set and R^2^ as the scoring metric (GridSearchCV function of the scikit-learn library in Python): constant that multiplies the L1 term in LASSO (1e-4, 1e-3, 1e-2, 1e-1, 1), the number of features to consider when looking for the best split in RF (square root, log2, 16%, 32%, 64%, 100% of all features), the maximum depth of the tree in RF (2, 4, 8, 16), the minium number of samples required to split an internal node in RF (2, 4, 8, 16), and the minimum number of samples required to be at a leaf node (1, 2, 4). We fixed the number of trees in RF model to 2,000.

## Data availability

All data (including metadata, sequencing data and metablomics data) have been deposited in the NCBI SRA under accession number PRJNA754019.

## Code availability

The customized scripts used in this study are available at: https://github.com/liaochen1988/DFdynamics.

## Acknowledgments

We would like to thank members of LD lab for constructive comments on the manuscript. This research was supported by National Key R&D Program of China (No.2019YFA0906700, to L.D.), National Natural Science Foundation of China (No.31971513, No. 32061143023, to L.D.) and China Postdoctoral Science Foundation (2020M682968, to H.L.).

## Author Contributions

H.L. and L.D. conceived the study. H.L. performed the experiments and analyzed the data. C. Liao analyzed the sequencing data and performed the computational analysis. J.T., J.C., C. Lei, L.Z. and L.W. assisted in experiments and/or data analysis. C. Liao, H.L. and L.D. wrote the manuscript with contributions from all coauthors.

