## Supplementary Figures for "Ecological dynamics of the gut microbiome in response to dietary fiber"

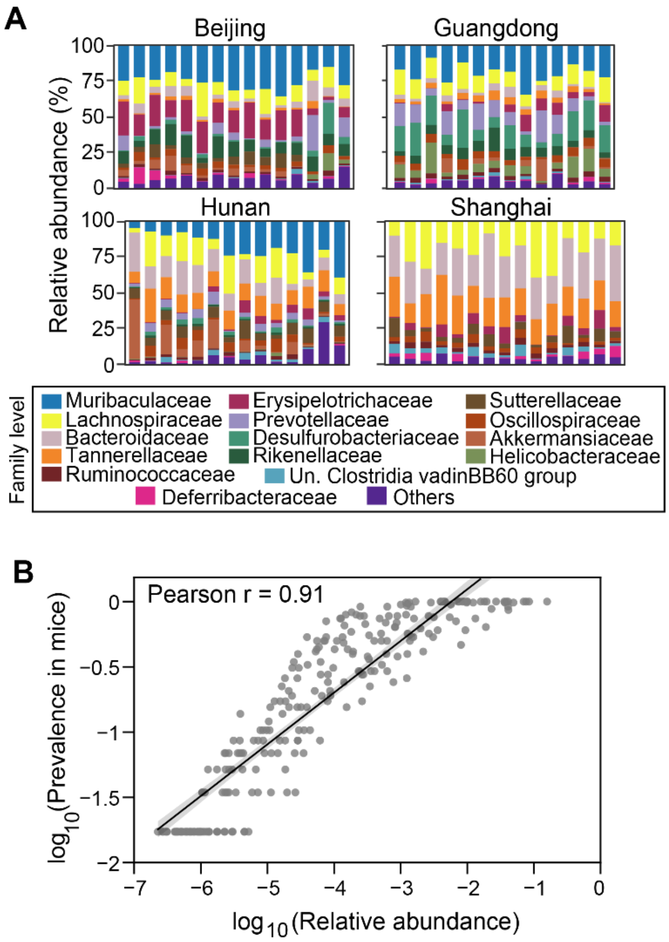


**Figure S1. Baseline gut microbiota. A.** Composition of the four vendors (Beijing, Guangdong, Hunan, Shanghai) at the family level. Bars represent individual mice. Adonis test indicates significant difference in the baseline gut microbiota composition across the four vendors (P<0.001). **B.** Linear relationship between relative abundance and prevalence (across individual mice from all vendors) of bacterial taxa grouped at the lowest taxonomic level (grey dots). Line: linear fit; shading area: 95% confidence interval.

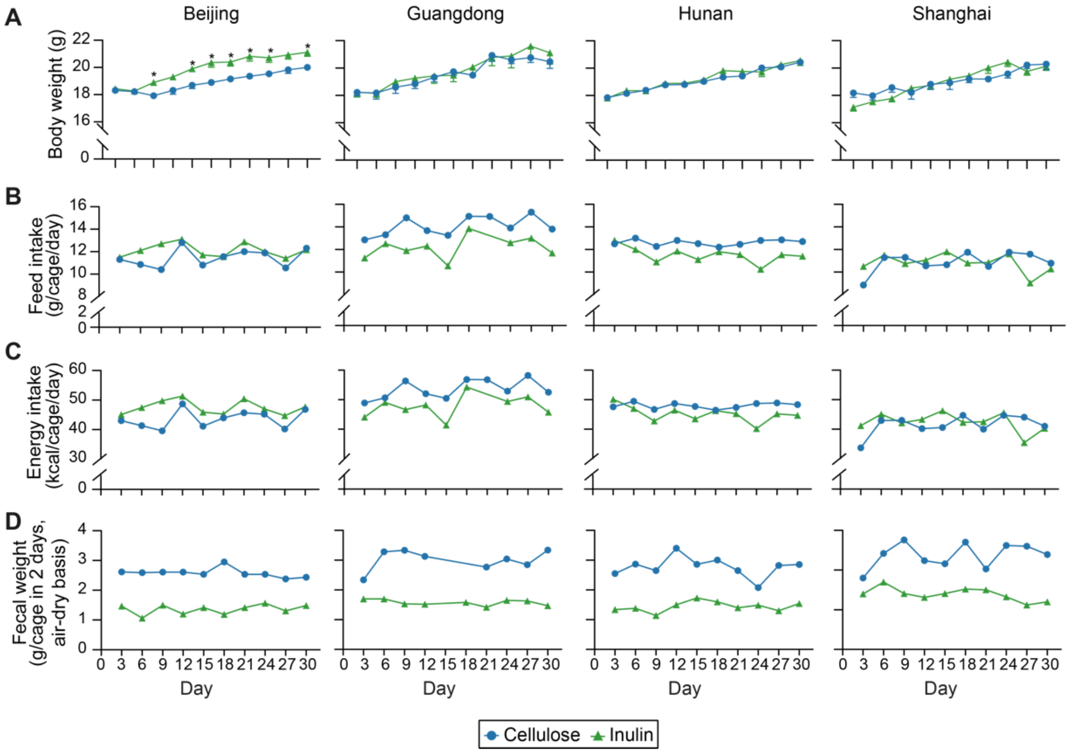


**Figure S2. Effects of inulin on (A) body weight, (B) daily food intake, (C) daily energy intake, and (D) 48-hr fecal sample weight of mice receiving inulin or cellulose supplementation.** Each symbol represents the mean body weight in panel A (bars: standard error of the mean; n=4 for Hunan and Guangdong, n=5 for Beijing and Shanghai) or a single data point in panels B-D (mice from the same vendor were co-housed). All food intakes were converted to energy intakes by multiplying food weight and its energy density (3.8 and 3.9 kcal/g for the cellulose- and inulin-based diets, respectively). The body weight data were analyzed by ordinary one-way ANOVA (Analysis of variance) with Turkey post hoc test between inulin and cellulose group. * $P < 0.05$.

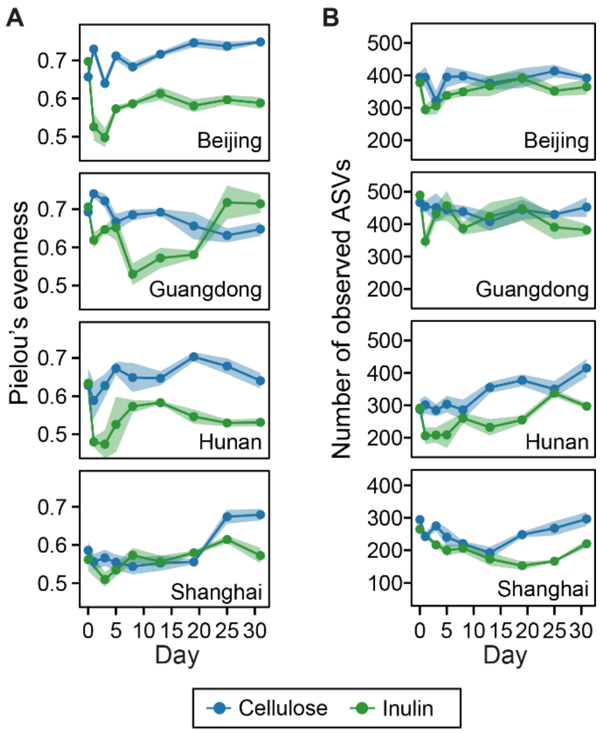


**Figure S3. Dynamics of (A) species evenness (Pielou's evenness) and (B) richness (number of observed ASVs) following inulin intervention.** Lines represent mean values across mice within the same vendor and shading areas represent standard error of the mean (n=4 for Hunan and Guangdong, n=5 for Beijing and Shanghai).

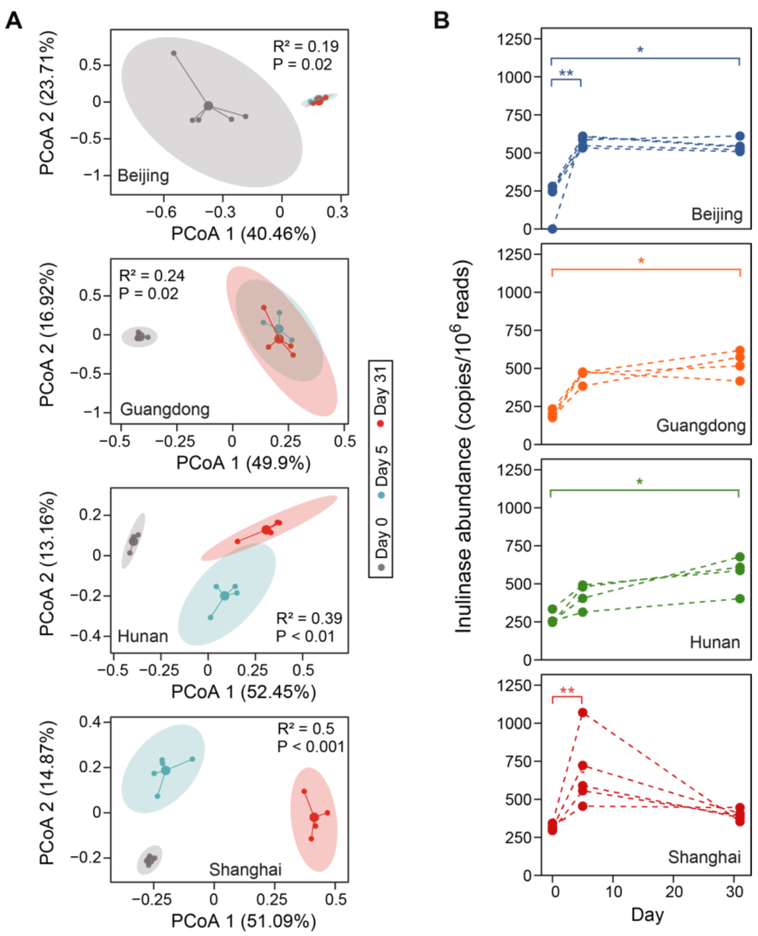


**Figure S4.** Relative abundance of gut microbiome genes following inulin intervention. **A**. High-dimensional gene family composition represented by robust PCoA (principal coordinate analysis) plot. Samples cluster by the day of collection and, for each cluster, smaller dots represent individual mice and the single bigger dot represents the cluster center. An eclipse was drawn around the cluster center to show its 95% confidence interval. $R^2$ and P-value were obtained from Adonis analysis, which tests for the difference in gene abundances among three representative timepoints during intervention (day 0: baseline, day 5: short-term response, day 31: long-term response). **B**. Relative abundance of inulinases/fructanases, calculated as the sum of reads mapping to individual CAZy genes (GH32, GH91 and CBM38) [1]. Each dotted line represents an individual mouse. *: P < 0.05; **: P < 0.01; ***: P < 0.001.

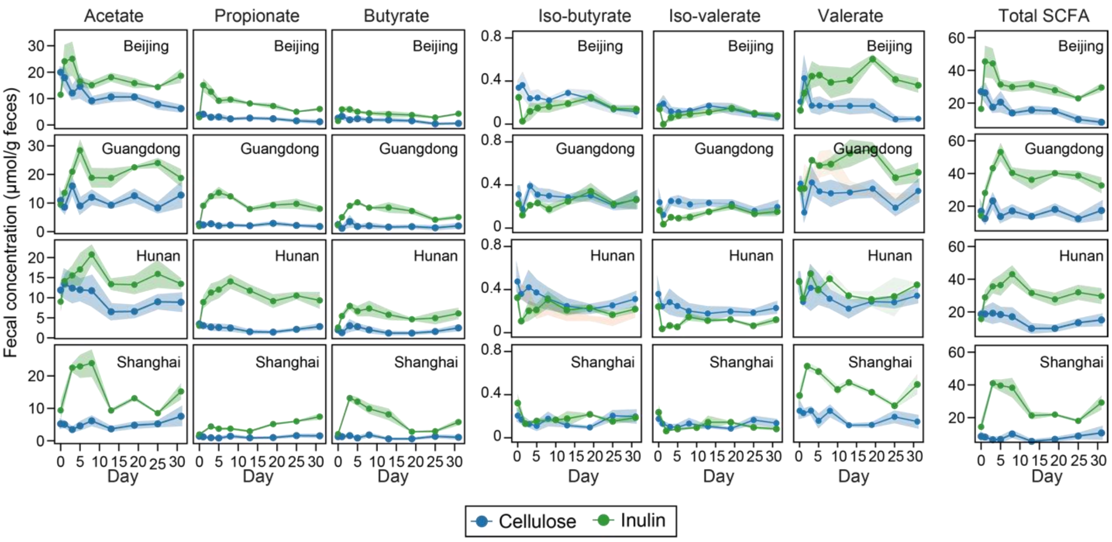


**Figure S5. Dynamics of fecal short-chain fatty acids (SCFAs) concentration following inulin intervention.** Total SCFAs include acetate, propionate, butyrate, iso-butyrate, iso-valerate and valerate. Dots/lines represent mean concentrations across mice from the same vendor and shading areas represent standard error of the mean (n=4 for Hunan and Guangdong, n=5 for Beijing and Shanghai).

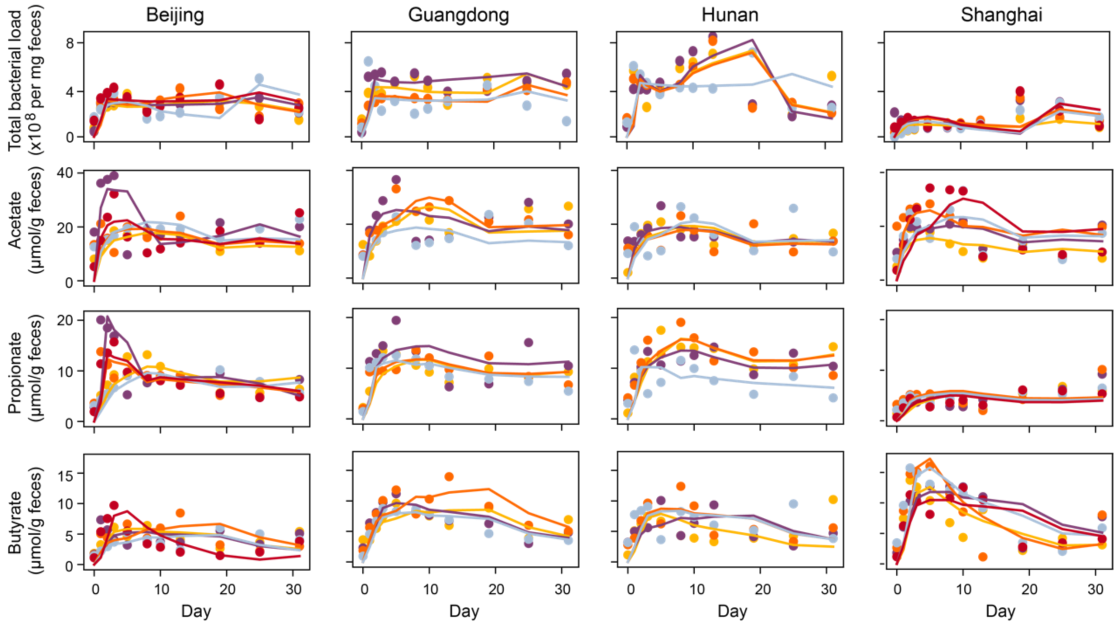


**Figure S6. Reconstructed time series (lines) of total bacterial load and three major short-chain fatty acids (acetate, propionate, butyrate) concentration by sequential non-negative matrix factorization (sNMF).** Dots represent observations, i.e., the original time series data from which reconstructed time series were built, and lines represent the reconstructed time series using the first two factors decomposed by sNMF. Lines and dots are color-coded on a per-mouse basis (n=4 for Hunan and Guangdong, n=5 for Beijing and Shanghai).

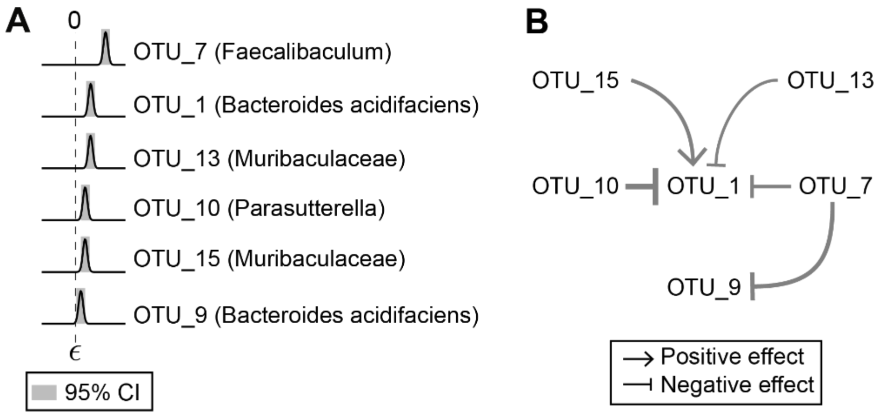


**Figure S7. Inferred ecological network of OTUs (Operational Taxonomic Units; 97% sequence similarity). A.** Posterior distribution of $\epsilon$ (the impact of dietary fiber on bacterial growth) for six putative primary degraders at the OTU level. OTUs are ranked according to their posterior mean of $\epsilon$. The shading area represent 95% credible interval (CI) of $\epsilon$. **B**. Ecological interactions among the six OTUs shown in the panel A. Point and blunt arrows represent positive and negative interactions respectively. The arrow thickness is proportional to the posterior mean of the corresponding interaction coefficient.

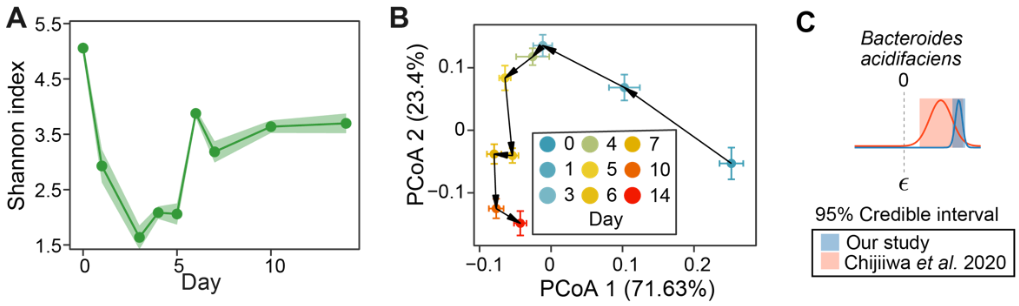


**Figure S8. Microbiome compositional analysis of previously published data from Chijiiwa *et al.* (n=9)** [2]**.** In this study, the shift in murine gut microbiota was tracked for two weeks following inulin intervention. **A**. Temporal shifts in alpha diversity of gut microbiota. **B**. Trajectory of gut microbiota composition shown in robust PCoA (principal coordinate analysis) plot. Each dot represents the mean principal coordinate score across all mice and the corresponding error bar represents the standard error of the mean. **C**. Posterior distribution of $\epsilon$ (the impact of dietary fiber on bacterial growth) for *Bacteroides acidifaciens* (the only taxa whose $\epsilon$ is positive and significantly different than zero).

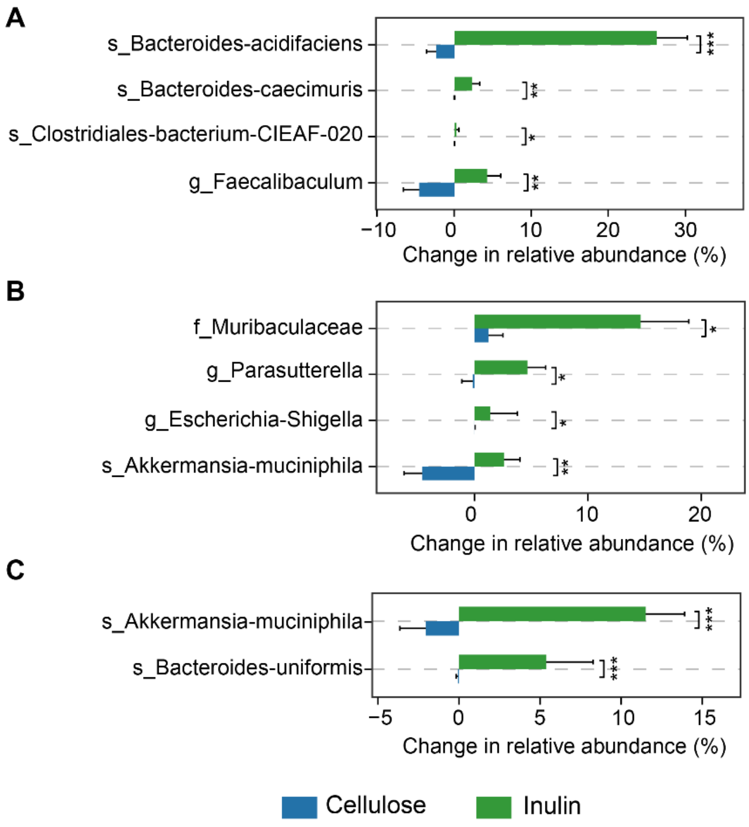


**Figure S9. Bacterial taxa with significant difference in relative abundance between the inulin group and the cellulose group.** Relative abundance changes were calculated between day 0 and day 1 (A), day 0 and day 5 (B), day 0 and day 31 (C). The bars represent the changes in the relative abundance of specified taxa across all mice from all vendors in the inulin or cellulose group. The error bars represent the standard error of the mean (n=18 for inulin and n=20 for cellulose). *P*-values were obtained from Wilcoxon rank-sum test after multiple test correction via false discovery rate (FDR) estimation. *, FDR < 0.05; **, FDR < 0.01; ***, FDR < 0.001.

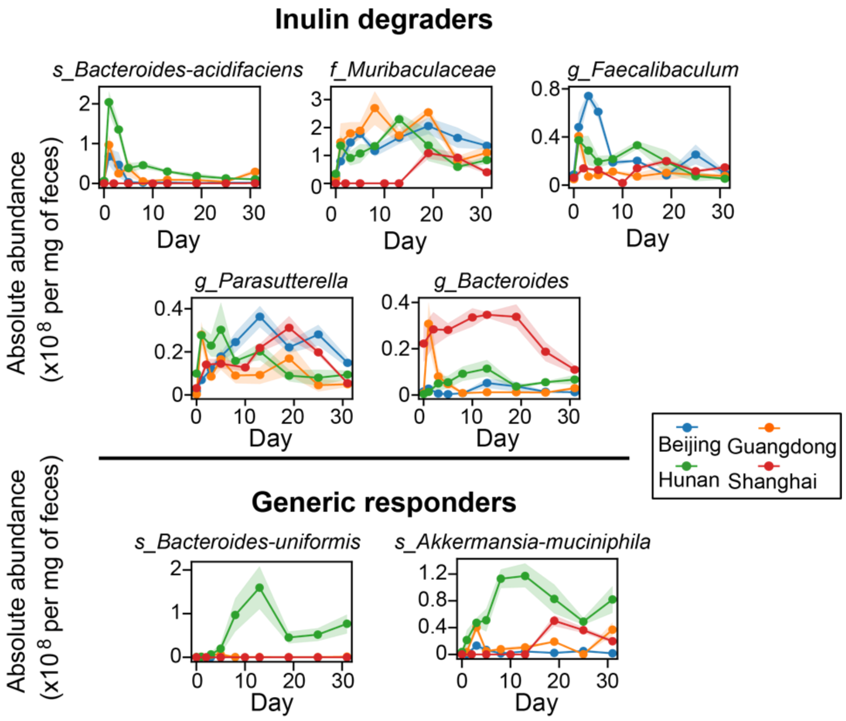


**Figure S10. Dynamical responses of the five inulin degraders and two generic responders.** Lines/dots: absolute abundance averaged across mice from the same vendor (n=4 for Hunan and Guangdong, n=5 for Beijing and Shanghai). Shading area: standard error of the mean values.

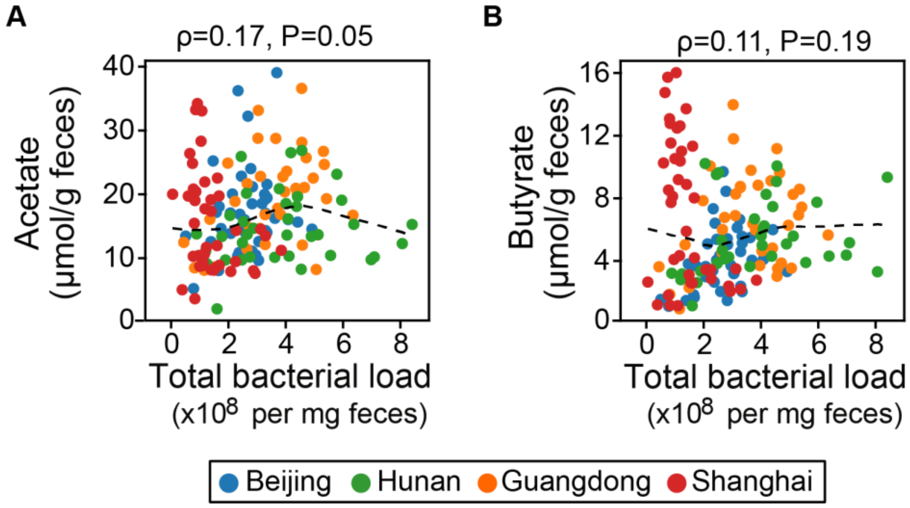


**Figure S11. Spearman correlation of total bacterial load with acetate (A) and butyrate (B) correlation.** Dots of the same color represent all samples from mice of the same vendor but collected at different timepoints. Dashed line: Lowess (Locally Weighted Scatterplot Smoothing) regression. Spearman correlation coefficient (ρ) and adjusted P-value (after Benjamini-Hochberg correction) are indicated in each plot.

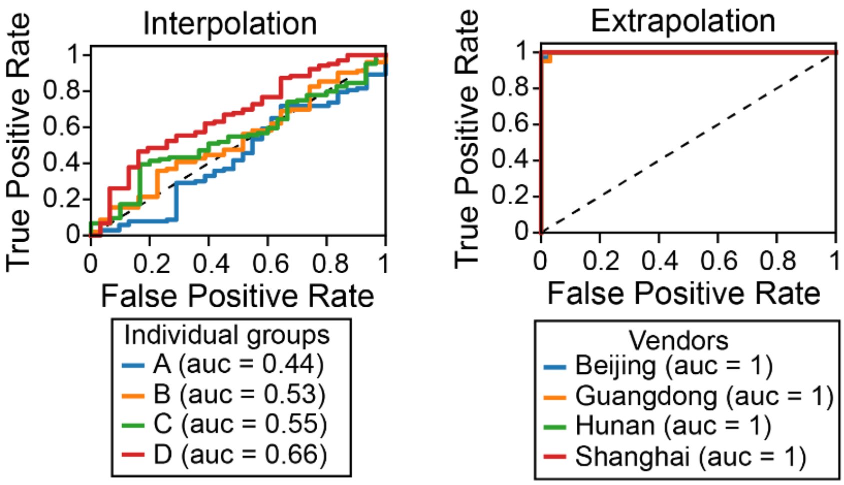


**Figure S12. Receiver operating characteristic (ROC) curve analysis of the similarity between training and testing datasets.** A Random Forest (RF) classifier trained to discriminate the two datasets outputs area under the ROC curve (AUC) as a similarity score. ROC curves were obtained by computing the probability of samples in the full datasets (both training and test sets) predicted as being taken from the training distribution. Specifically, we first concatenated the training and test sets and assign labels 1 and 0 respectively. The new combined dataset was then stratified into 20 folds and each time, a RF classifier was trained on 19 folds and the used to predict the probability of the remaining fold being sampled from the training set. See **Methods** of the main text for details of the two data-split strategies ("interpolation" and "extrapolation"). When the full dataset was split by the "interpolation" approach, training data and test data are nearly indistinguishable from each other (AUC close to 0.5); in contrast, for "extrapolation", training data and test data are fully distinguishable from each other (AUC close to 1).

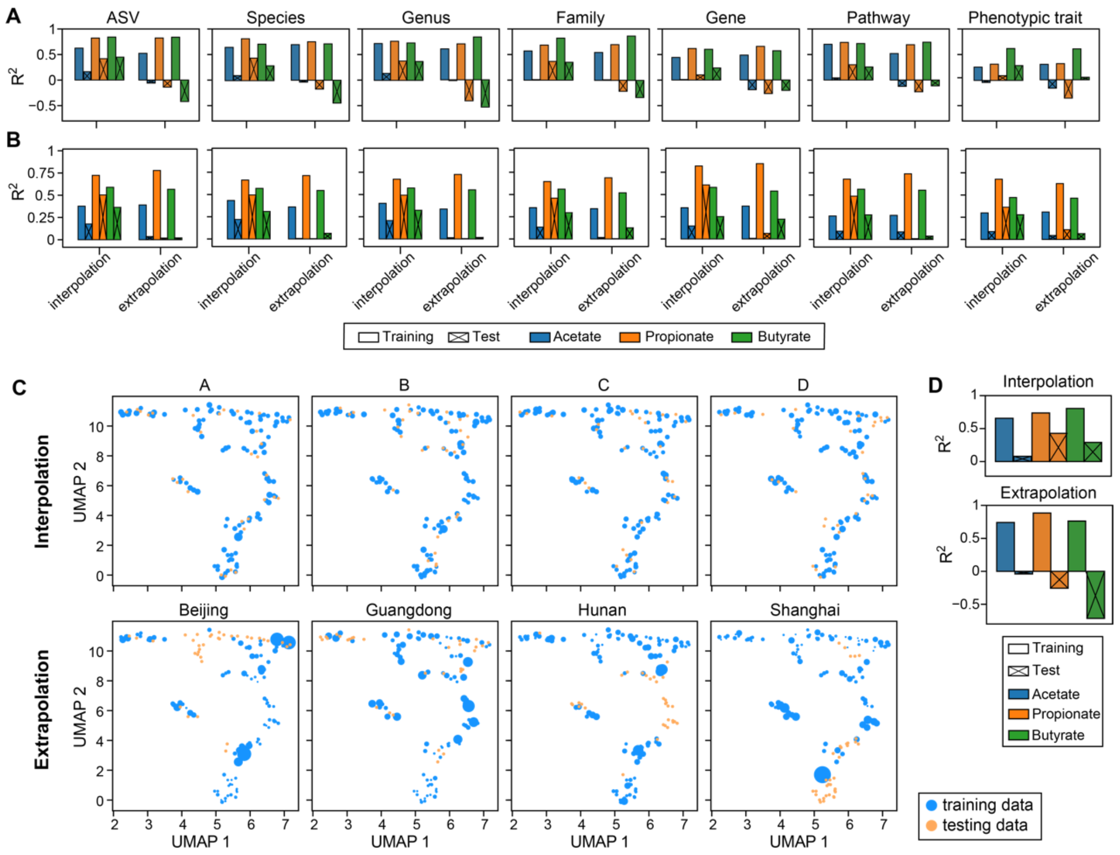


**Figure S13. Poor performance of Random Forest (RF) regression model in predicting short-chain fatty acids (SCFAs) concentration (see Fig. 5C of the main text for the results) cannot be rescued by using (A) alternative predictors, (B) alternative regression models, and (C,D) weighting of training samples. A**. Prediction accuracy of a RF model trained on different taxonomic- (ASV, Species, Genus, Family) or functional- (Gene, Pathway, Phenotypic trait) predictors. Phenotypic traits include 41 binary variables that mainly describe the capability of bacteria to utilize sugars and to de novo synthesize amino acids (prototrophy vs. auxotrophy). The abundances of genes, pathways and phenotypic traits were predicted using PICRUSt2 [3]. **B**. Prediction accuracy of the MelonnPan algorithm [4] trained on the same predictors as used in panel A. Notably, MelonnPan predicts relative profiles of SCFAs from relative abundance of gut microbiota. **C**. Weights assigned to the training data. The gut microbiota composition of all samples was shown in a reduced two-dimensional UMAP (Uniform Manifold Approximation and Projection) space [5]. The bigger the weights, the larger circle sizes. Following the same approach as described in the legend of **Fig. S12**, we obtained the probability ($p_i$) of each sample in the full dataset being predicted as taken from the training subset. The weight assigned to sample $i$ was then given by $(1 - p_i)/p_i$, which makes intuitive sense: the higher the numerator and the lower the denominator, the closer the sample gets to the high-density regions of the test data. **D**. Prediction accuracy of an RF model built from weighted training data.

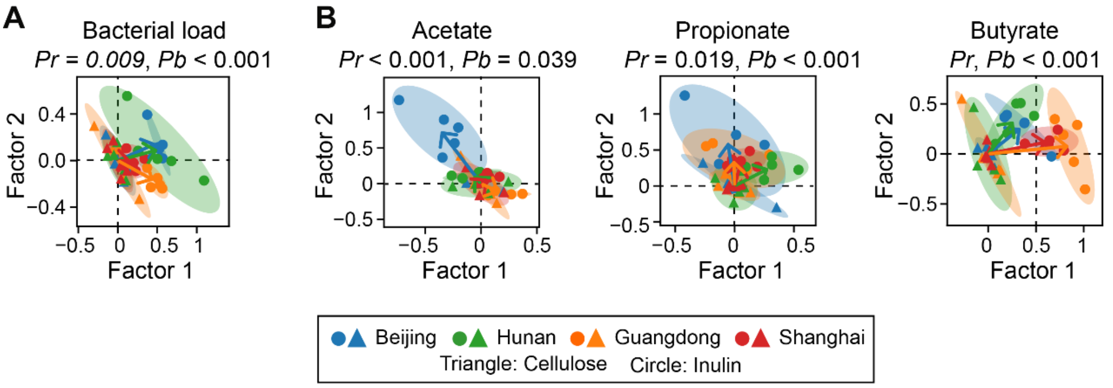


**Figure S14. Reduced 2-dimensional representation of the resistant starch-induced responses in bacterial load (A) and three major SCFAs (B).** The same figure legend applies as in the main text **Fig. 3B, C**.

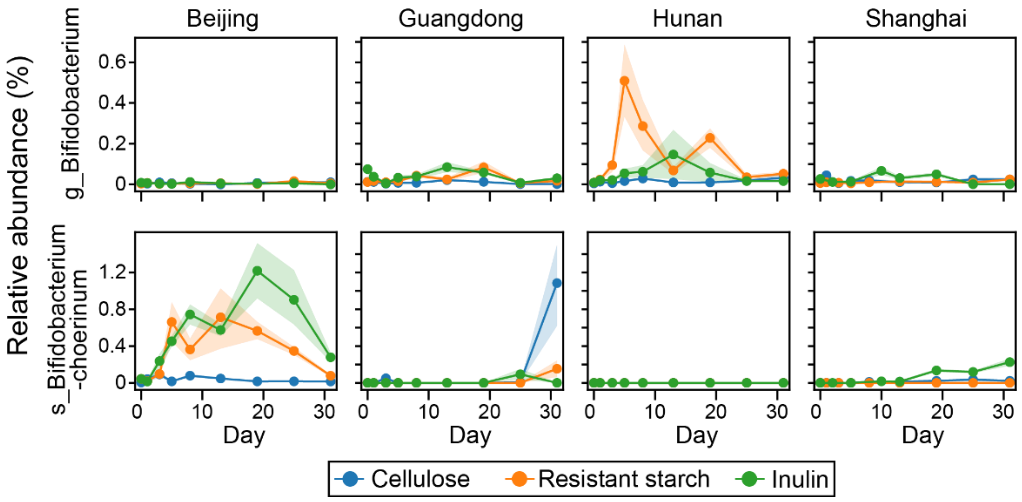


**Figure S15. Temporal shifts in the relative abundance of Bifidobacterium-related taxa following inulin and resistant starch intervention.** The two taxa, g_Bifidobacterium and s_Bifidobacterium-choerinum, are present (using a relative abundance threshold of 1e-5) in 17% and 66% of all baseline samples (regardless of vendor and diet) respectively. Specifically, g_Bifidobacterium is consistently present in >50% of the baseline samples from each vendor. s_Bifidobacterium-choerinum is present in some baseline samples of Beijing and Shanghai mice but completely missing (i.e., relative abundance equals to 0) in all baseline samples from Hunan mice.

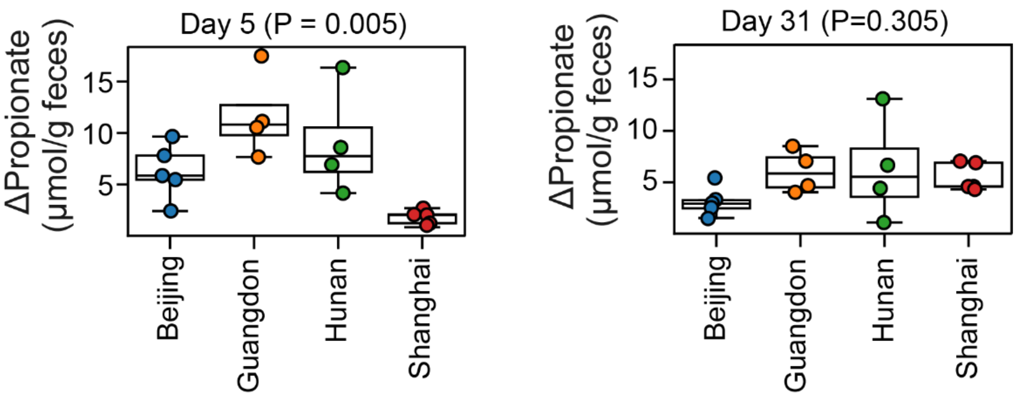


**Figure S16. The between-vendor difference of pre-to-post changes in propionate concentration (Δpropionate = endpoint value - baseline value) using day 5 (left panel) or day 31 (right panel) as the intervention endpoint.** The P-values were obtained by Permutational Multivariate Analaysis of Variance (PERMANOVA) with Minkowski distance as the distance metric**.** Each dot represents a mouse and dots of the same color represent mice from the same vendor (n=4 for Hunan and Guangdong, n=5 for Beijing and Shanghai).

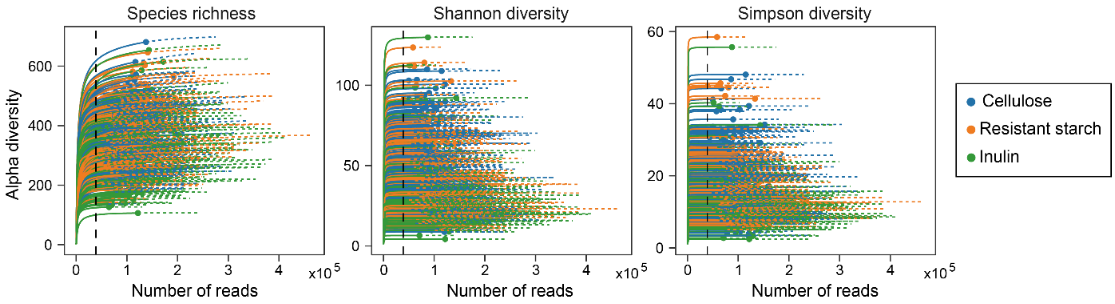


**Figure S17. Rarefaction analysis of 16S rRNA amplicon sequencing data.** Rarefaction curves were generated using the iNEXT package[6]. Solid lines represent the observed alpha diversity with the number of reads sampled, and dashed lines represent the extrapolation of the solid lines until 25% more reads. To avoid sample-to-sample bias due to variable sequencing depth (different number of reads per sample), all samples were rarefied to 38,980 sequences (black dashed line) per sample before downstream analysis.
